# Post-transcriptional control of homeostatic B-cell signalling by HuR is required for innate B cell maintenance and function

**DOI:** 10.1101/2024.09.17.611791

**Authors:** Dunja Capitan-Sobrino, Mailys Mouysset, Orlane Maloudi, Yann Aubert, Ines C. Osma-Garcia, Maia Nestor-Martin, Trang-My M. Nguyen, Greta Dunga, Manuel D. Diaz-Muñoz

## Abstract

Innate B-1 cells constitute a self-maintained layer of defence for early detection of bacteria, clearance of apoptotic cell debris and removal of autoantigens driving autoimmunity. B-1 cells are originated from foetal tissues but, as opposed to B-2 cells, the molecular mechanisms behind their development and homeostatic maintenance remain largely unknown. Here we demonstrate that post-transcriptional regulation by the RNA binding protein HuR is essential for the homeostatic self-replenishment of innate B-1 cells, the expansion of B-1 cell clones targeting self-antigens and the production of natural autoantibodies. HuR KO B-1 cells fail to express the high levels of surface BCR, TACI and BAFFR required for tonic signalling and cell survival. Mechanistically, HuR binds to the 3’UTRs of mRNAs encoding these surface receptors and of pro-survival molecules, like BCL-2 and MCL-1, promoting their translation into protein. In summary, we reveal the need of post-transcriptional regulation in BCR expression, tonic signalling and homeostatic maintenance of functional B-1 cells.

## INTRODUCTION

B-1 cells constitute a unique subset of B cells with innate-like and regulatory properties. They provide a first line of defence against bacteria and self-antigens, differing from conventional B-2 cells in ontogeny, phenotype and function. Early characterization of B-1 cells defined them as long-lived, self-renewing cells with a restricted immunoglobulin repertoire biased to the production of poorly mutated IgM autoantibodies ^1, 2, 3^. Expression of the scavenger receptor CD5 classify B-1 cells into two major subsets. CD5^+^ B-1a cells produce the vast majority of IL-10 and autoantibodies while CD5^-^ B-1b cells exert mixed innate-like and adaptive functions. B-1 cells mainly populate the pleural and peritoneal cavity in mice. However, they also reside in mucosal and non-mucosal tissues (gut, skin, lung, kidney) to orchestrate early immune responses against bacteria through the polarization of macrophages, and the production of antibodies and cytokines ^4, 5, 6^. This is largely possible due to the overrepresentation of B-1 cells with rearranged V_H_11 and V_H_12 gene segments ^3, 7^, which encode antibodies against phosphatidylcholine (PtC), a major lipid exposed in mucosal tissues, apoptotic cell bodies and bacteria. These PtC-specific B-1a cells emerge with age in a process of immunodominance against commensal bacteria-derived antigens ^1, 8^ and, possibly, self-dead cells. In adult mice, PtC-reactive B-1a cells can account for up to 50% of total B-1a cells in the peritoneal cavity, in deep contrast to the polyclonal repertoire found in neonates ^9^. In humans, monoclonal expansion of mature CD5^+^ B-1a cells can lead to B-cell chronic lymphocytic leukaemia (B-CLL) while alterations in self-antigen detection and/or in the regulatory function of B-1a cells, e.g. through the provision of IL-10, are often found in autoimmune diseases such as arthritis and lupus ^10, 11^.

B-1 cell progenitors emerge early during development in the yolk sac (YS) and splanchnopleura in a hematopoietic stem cell (HSC)-independent manner ^12, 13^. Then, following a second wave of development, they migrate into the foetal liver and spleen in which premature recombination of the Ig kappa gene allows assembly of a mature B-cell receptor (BCR), bypassing negative B cell selection and generation of autoreactive B-1 cells ^14, 15^. In neonates, terminal B-1 development takes place in liver and spleen before the establishment of the mature B-1 cell compartment in body cavities from day 7 after birth ^16^. Contrary to B-2 cells, bone marrow transfer has limited capacity to regenerate B-1 cells in adult mice, suggesting that the expansion and maintenance of this cell population is mainly mediated by selection and self-renewal ^17^. Alternatively, recent cell barcoding experiments in transgenic mice highlight the possibility of the presence of a common progenitor that can give rise to both B-2 and B-1 cells ^18, 19^. However, the extension of this crosstalk in adults might be limited by the presence of mature B-1 cells that interfere with the establishment of newly developed B-1 cells, possibly through a self-antigen mediated competition mechanism ^18, 20^.

Identification of unique transcriptional and post-transcriptional master regulators of B-1 cell development, maintenance and function remains elusive. It is somehow expected that important transcriptional factors (TFs) in B-2 cell development, like PAX-5, EBF1, E2A and IKAROS ^21^, might also control B-1 cell lymphopoiesis. However, recent evidence showing the negative regulatory function of the TF IKAROS in B-1 cell compartment generation highlights the importance of dedicated studies in B-1 cells ^22^. In this line, it has been demonstrated that the LIN28-Let7a axis acts as a post-transcriptional switch that controls the expression of ARID3A for foetal lymphopoiesis ^23^, selection and establishment of the autoreactive B-1 cell pool ^24, 25^. The TFs BHLHE41, TCF1 and LEF1 are likely to play an important role during peripheral B-1 cell renewal and long-lived maintenance by keeping in check exacerbated BCR signalling and downstream proliferative programs ^26^. Instructive BCR signalling shapes the B-1 cell repertoire during self-renewal ^27^ and, along with the cytokine receptors TACI, BAFFR and IL5R, shapes the long-term maintenance and function of B-1 cells ^28^. In this study, we uncover the RNA binding protein (RBP) HuR as a key post-transcriptional regulator controlling the expression of the BCR and pro-survival receptors TACI and BAFFR for B-1 cell self-replenishment and expansion during T-independent type II immunity.

Post-transcriptional control of RNA constitutes a complex, but essential, layer of regulation between the early event of gene transcription and final protein synthesis ^29, 30^. RBPs and non-coding RNAs (ncRNAs) cooperate or compete during the regulation of RNA splicing, editing, stability, location and translation. Highly specialised molecular machineries segregate these processes in time and space while multifunctional RBPs, such as HuR, have a prevalent role on finely tuning each of these processes in an RNA target-specific manner. Nowadays, a handful of RBPs have been studied mostly in B-2 cells. They have irreplaceable functions from progenitor commitment into the B-cell lineage to the establishment of a durable immune response. Splicing factors such as PTBP1, TIA1 and TIAL1 control B-cell reprograming upon activation, promote the expression of the DNA repair machinery and the expansion of antigen-specific B cell clones ^31, 32, 33^. RBPs of the exosome and RNA decay complexes essentially contribute to the diversification of B-cell receptors during development ^34, 35^. RNA surveillance and timely regulation of messenger (m)RNA stability coordinate B cell selection, proliferation and differentiation both during B-cell development, maturation and terminal differentiation into antibody-secreting cells ^36, 37, 38^. Here we demonstrate that, while dispensable during B-1-cell development, the RBP HuR is essential for the self-maintenance of mature B-1 cells and the expansion of autoreactive B cell clones for the clearance of apoptotic cells.

HuR is the only member of the *Elavl-* family of proteins expressed in immune cells. It is a multifunctional RBP that acts as splicing regulator in the nucleus or as a modulator of mRNA stability and translation in the cytoplasm. HuR contains three RNA recognition motifs through which it targets AU-and U-rich regulatory elements present close to the 3’ splice site of unprocessed introns or within 3’ untranslated regions (3’UTR) of mature mRNAs ^39, 40^. HuR is ubiquitously expressed in all cells. However, post-translational modification of HuR upon cellular sensing of external and internal cues control its protein stability, subcellular localization and RNA target specificity and affinity ^41^, leading to the regulation of different processes in the same cell type depending on its cellular status. This is exemplified in B cells in which HuR not only promotes the splicing of *Dlst*, one of the three components of the α-ketoglutarate dehydrogenase complex, for energy fuelling and scavenging of reactive oxygen species (ROS) upon activation of naïve B cells ^40^, but also controls affinity maturation of already activated germinal centre (GC) B cells by controlling Myc mRNA translation and cell cycle progression ^42^.

In this study, we have combined the use of conditional knock out mouse models, molecular mapping of the HuR:RNA interactome and interrogation of the cell transcriptome to uncover the unique functions of HuR in B-1 cells. The high expression of HuR in B-1 cells contributes to the mRNA translation of the cell surface receptors BCR, TACI and BAFFR for homeostatic survival, self-renewal, and production of natural autoantibodies. In addition, by using a type II immunization model with thymic-derived apoptotic bodies, we demonstrate that HuR promotes the expansion of self-reactive B-1 cell clones for the clearance of apoptotic cell debris.

## RESULTS

### Innate B-1 cells are reduced in the absence of HuR

Comparative analysis of the expression of HuR in different B cell subsets revealed a 1.7-fold increase in protein levels in innate B-1 cells compared to B-2 cells (***Fig 1A***). This increment was very consistent across experiments and could be observed in B-1 cells from the peritoneal cavity and the spleen (***Suppl. Fig. 1A***). Importantly, HuR mRNA abundance remained unaltered in both cell types (***Fig 1B***) suggesting the existence of a cell-intrinsic post-transcriptional mechanism that promotes high expression of HuR protein in B-1 cells.

**Figure 1.**
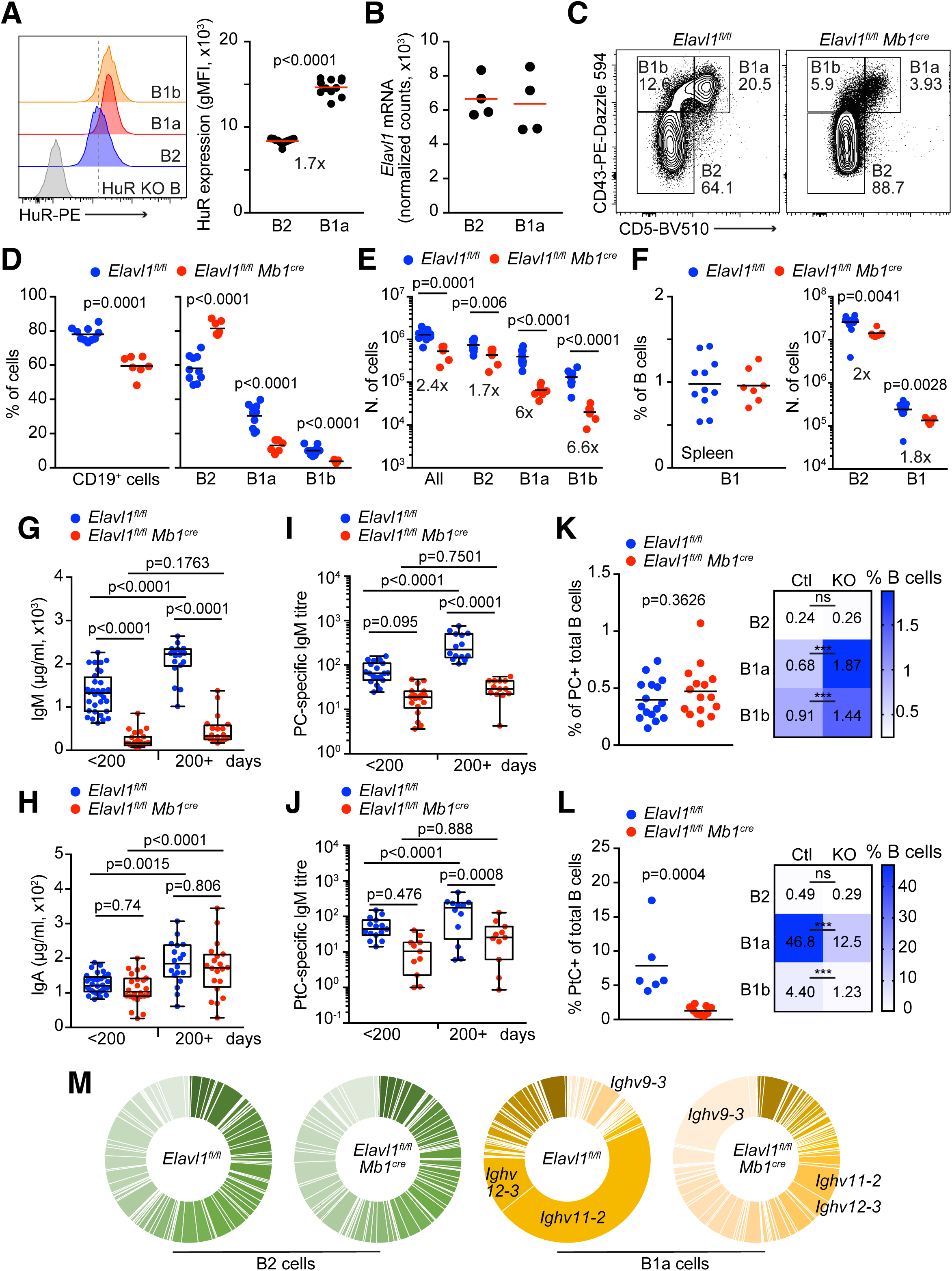
HuR is required for innate B-1 cell maintenance and natural autoantibody production. **A,** Analysis of HuR expression in B-1 and B-2 cells from the peritoneal cavity. Left panel, representative histogram showing HuR protein levels. Right panel, quantitation of HuR geometric mean fluorescent intensity (gMFI) detected by flow cytometry. Data pooled from two out of more than three independent experiments with a minimum of 3 mice. **B,** Quantitation of *Elavl1* mRNA abundance in peritoneal cavity B-1 and B-2 cells measured by mRNAseq (n=4 biological replicates per genotype). **C,** Contour plots showing the percentage of CD19^+^ B-1a, B-1b and B-2 cells present in the peritoneal cavity of control and *Elavl1^fl/fl^ Mb1^Cre^* mice. Data representative from more than three independent experiments. **D, E,** Percentage and number of total CD19^+^ B cells, and B-1a, B-1b and B-2 cells found in the peritoneal cavity of control and *Elavl1^fl/fl^ Mb1^Cre^*mice. Data pooled from two out of more than three independent experiments performed with a minimum of 3 mice per genotype. **F,** Percentage and number of splenic B-1 and B-2 cells in the same mice showed in D, E. **G, H,** Quantitation of the IgM and IgA levels found in the serum of young (<200 days) and old (>200 days) control and *Elavl1^fl/fl^ Mb1^Cre^* mice. Data is from a minimum of 18 mice per genotype and age collected in at least three independent experiments. **I, J,** Measurement of natural IgM antibodies against the self-antigens PC and PtC in young and old control and *Elavl1^fl/fl^ Mb1^Cre^* mice. Data is from a minimum of 6 mice per genotype and age collected in two (J) or three (I) independent experiments. **K, L,** Percentage of PC- and PtC-specific B cells detected in the peritoneal cavity of control and *Elavl1^fl/fl^ Mb1^Cre^* female mice by flow cytometry (mouse age > 200 days). Right panel, heatmaps showing the distribution of PC- and PtC-specific B cells between the B-1a, B-1b and B-2 cell subsets of the peritoneal compartment (*** p<0.001). Data is from the same mice shown in I, J. **M,** Pie charts representing the frequency of *Ighv* gene expression in B-2 and B-1a cells isolated from the peritoneal cavity of control and *Elavl1^fl/fl^ Mb1^Cre^* mice (mRNAsq data, n=4 replicates per genotype). Two-tailed Mann-Whitney tests were used in A, D, F, K and L. One-way ANOVA followed by multiple t tests were performed in D, E, G, H, I, J, K and L.

To uncover the importance of HuR in B-1 cells, we generated B-cell conditional knock out (KO) mice (*Elavl1^fl/fl^ Mb1^Cre^* ^40, 43^) and quantified by flow cytometry the percentage and number of B-1 and B-2 cells in the peritoneal cavity and spleen of these mice and their littermate controls (***Fig 1C-F*** *and* ***Suppl. Fig. 1B*** and ***1C***). Overall, we found the proportion and number of total CD19^+^ cells to be significantly decreased in *Elavl1^fl/fl^ Mb1^Cre^* mice (***Fig. 1D***, ***1E*** and ***Suppl. Fig. 1D***). Segregation by cell type revealed that B-2 cell numbers were mildly impacted in the absence of HuR, with a reduction between 1.7 to 2-fold depending on the compartment analysed (***Fig. 1E*** and ***1F***). By contrast, B-1a and B-1b cells were far more affected by the absence of HuR. The proportion and number of B-1a and B-1b cells were decreased by more than 6-fold in the peritoneal cavity of *Elavl1^fl/fl^ Mb1^Cre^* mice when compared to control mice (***Fig 1C-E***). The percentage of B-1 cells in the spleen was similar in control and *Elavl1^fl/fl^ Mb1^Cre^* mice (***Fig 1F*** *and* ***Suppl. Fig. 1D***). However, the total number of splenic B-1 cells was reduced by 1.8-fold in *Elavl1^fl/fl^ Mb1^Cre^* mice (***Fig 1F***). This decrease was in line with the overall reduction of the B-cell compartment observed in the spleen of *Elavl1^fl/fl^ Mb1^Cre^* mice (***Suppl. Fig. 1D***). Altogether, our data suggest that HuR is required for the maintenance of innate B-1 cells in mice.

### Impaired natural autoantibody production in the absence of HuR

B-1 cells from body cavities are major producers of natural IgM autoantibodies and, in gastrointestinal lymphoid tissues and other mucosal sites, can efficiently undergo class-switch recombination (CSR) to produce IgA in response to food antigens and microbiota ^5^. In accordance with the low number of B-1 cells in *Elavl1^fl/fl^ Mb1^Cre^* mice, we observed a profound reduction in the total levels of IgM in the blood of these mice (***Fig 1G***). Control mice steadily increase their blood IgM levels as they aged. However, this increment was not observed in aged *Elavl1^fl/fl^ Mb1^Cre^* mice (***Fig 1G***). IgA levels were neither altered in young or aged *Elavl1^fl/fl^ Mb1^Cre^* mice (***Fig 1H***), suggesting that impaired B-1 cell numbers in these mice mostly impacts the production of natural IgM antibodies. To confirm this observation, we measured the antibody titres of two commonest natural IgM antibodies recognising the self-antigens phosphorylcholine (PC) and phosphatidylcholine (PtC) released upon cell death. PC- and PtC-specific IgM titres were reduced, although not significantly, in young *Elavl1^fl/fl^ Mb1^Cre^*mice when compared with their control littermates (***Fig 1I*** and ***1J***). Importantly, and in deep contrast to control mice, *Elavl1^fl/fl^ Mb1^Cre^* mice failed to significantly upregulate the production of PC- and PtC-specific IgM autoantibodies as they aged (***Fig 1I*** and ***1J***). Analysis of PC-specific B cells in the peritoneal cavity of aged mice revealed that the percentage of these cells remained constant in *Elavl1^fl/fl^ Mb1^Cre^* mice or slightly increased when considering only the major auto-antibody producers B-1a and B-1b cells (***Fig 1K***). This suggested that the overall reduction of the B-1 cell compartment in the peritoneal cavity was likely responsible for the impaired production of anti-PC IgM antibodies in *Elavl1^fl/fl^ Mb1^Cre^*mice, rather than changes in the proportions of PC-specific B cells. By contrast, the percentage of PtC-specific B cells was profoundly decreased in the peritoneal cavity of *Elavl1^fl/fl^ Mb1^Cre^* mice, both when considering total B cells or B-1 cell subsets only (***Fig 1L***). Thus, we conclude that HuR expression is needed for the maintenance of B-1 cell clones producing natural autoantibodies.

To assess the extension of the changes in the BCR repertoire of B1 cells in the absence of HuR, we measured by mRNAseq the abundance of each *Ighv* mRNA transcript in B-2 and B-1a cells isolated from control and *Elavl1^fl/fl^ Mb1^Cre^* mice. While the pool of B-2 cells maintained a polyclonal BCR repertoire, we observed extensive changes in *Ighv* gene usage in HuR KO B-1a cells compared to control cells (***Fig 1M***). B-1a cells progressively restrict their BCR repertoire expanding those that recognise the most common self-antigens ^14^. Indeed, control B-1a cells showed an oligoclonal repertoire in which transcripts from the genes *Ighv11-2* and *Ighv12-3,* that encode BCR-isoforms specific against PtC, were overrepresented (***Fig 1M***). By contrast, HuR KO B-1a cells maintained a polyclonal repertoire closer to the one found in neonates ^9^. This was exemplified by a 7-fold and 3.5-fold reduction in the frequency of *Ighv11-2* and *Ighv12-3* genes. By contrast, the commonly low-abundant gene *Ighv9-3* was increased by almost 5-fold in HuR KO B-1a cells, although its antigen specificity remains unknown. Taken together, our data demonstrate that HuR contributes to the restrictive expansion of autoreactive B-1 cell clones.

### HuR is dispensable during innate B cell development

Innate B-1 cells follow a highly distinctive developmental pathway, with hematopoietic stem cell (HSC) progenitors first arising in the yolk sac (YS) and the para-aortic splanchnopleura (p-SP) for later differentiating into B-1 cell progenitors in foetal liver and spleen. Then, terminal differentiation takes place in the spleen of new-born mice, with mature B-1 cells homing into the body cavities after day 7 post-birth (***Fig. 2A***) ^16^. Embryonic B-1 cell progenitors express low levels of B220, and they can be tracked by the high-expression of CD19 and CD93 (***Fig. 2B***). To assess whether HuR was implicated in the early development of B-1 cell progenitors, we first confirmed efficient deletion of HuR in embryonic B cells from *Elavl1^fl/fl^ Mb1^Cre^*mice (***Fig. 2B***). Later quantitation of B-1 and B-2 progenitors in control and *Elavl1^fl/fl^ Mb1^Cre^* embryos (17.5-days of age, E17.5) revealed an increment in the percentage, but not in the numbers, of B-1 cell progenitors in the absence of HuR (***Fig. 2C***). Similarly, we found that the proportion and number of B-1 progenitors was augmented in the liver and spleen of *Elavl1^fl/fl^ Mb1^Cre^* new born mice (***Fig. 2D***). Thus, we conclude that HuR is not required for early B-1 cell development in the embryo and neonates.

**Figure 2.**
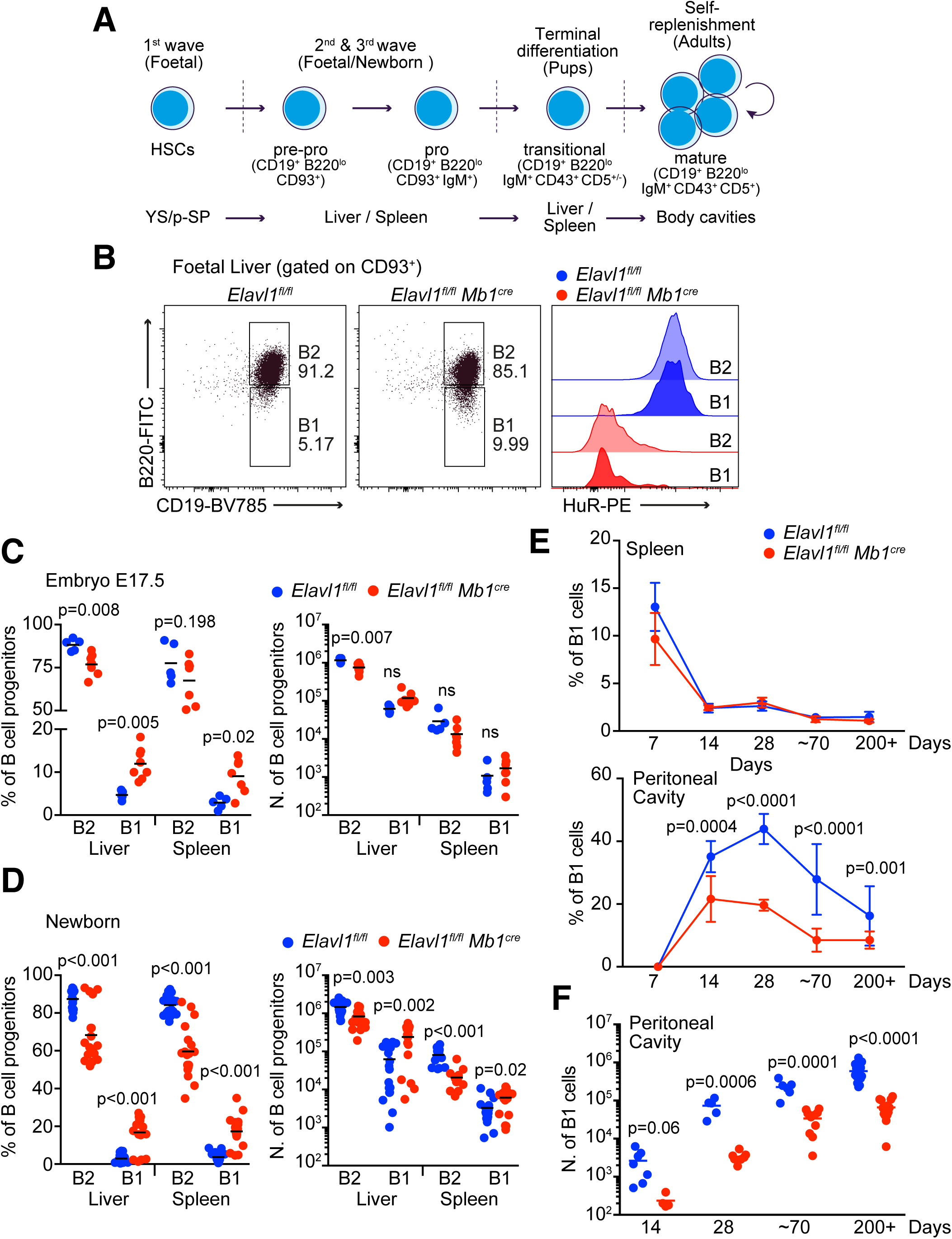
HuR is dispensable during innate B cell development. **A,** Graphical summary of B-1 cell development in mice. **B,** Analysis of efficient HuR deletion in foetal B cell progenitors. Left panels show representative FACs dot plots for quantitation of the proportion and number of B-1 and B-2 cell progenitors in foetal livers from control and *Elavl1^fl/fl^ Mb1^Cre^* mice. Right panel, representative histogram showing HuR expression in foetal B-1 and B-2 cell progenitors from these mice. **C,** Percentage and number of B cell progenitors in the liver of E17.5 embryos. Data from two independent experiments with a minimum number of 5 mice. **D,** Percentage and number of B cell progenitors in the liver and spleen of new-born mice. Data from three independent experiments performed with at least 7 mice. **E,** Proportion of B-1 cells found in the spleen (top panel) and peritoneal cavity (bottom panel) of control and *Elavl1^fl/fl^ Mb1^Cre^*mice at the indicated time points after birth. **F,** Quantitation of peritoneal B-1 cells found in mice shown in E. Data shown in E and F is from at least 5 mice per genotype collected in at least two independent experiments depending on the time point. Mouse genotype in C, D and E (2-weeks old or lower) was determined by intracellular flow cytometry analysis of HuR expression in B cell progenitors at the end of each experiment. Multiple two-tailed t tests were performed for statistical analysis of each of the panels.

Next, we questioned whether HuR was required for B-1 cell maturation and expansion of the B-1 cell compartment after birth. We measured the proportion and numbers of B-1 cells in the spleen and peritoneal cavity every 7 days after birth, at weaning (28 days), in young mice (∼70 days) and in aged mice (200+ days) (***Fig. 2E*** and ***2F***). Splenic B-1 cells represented a much larger population within B cells in young pups than in older pups and adults. In line with our previous observations (***Fig. 1F***), we found no differences in the proportion of splenic B-1 cells between control and *Elavl1^fl/fl^ Mb1^Cre^* mice at any of the time points analysed. B-1 cells did not populate the peritoneal cavity until between day 7 and day 14 after mouse birth (***Fig. 2E***). B-1 cells could be detected in the peritoneal cavity of both control and *Elavl1^fl/fl^ Mb1^Cre^* mice from day 14, indicating that HuR KO B-1 cells could migrate into body cavities from the spleen. However, the proportion and numbers of peritoneal B-1 cells were significantly reduced in *Elavl1^fl/fl^ Mb1^Cre^* mice from day 14 onwards (***Fig. 2E*** and ***2F***). Altogether, our data suggest that HuR is not required for B-1 cell maturation, but for the establishment and expansion of the B-1 cell compartment.

### HuR is required for the self-renewal and survival of innate B cells

In contrast to B-2 cells, which are constantly replenished from the bone marrow, B-1a cells are maintained in the adulthood by self-renewal. Indeed, analysis of cell cycling, by measuring the incorporation of the thymidine analogue BrdU, showed a 3-fold increase in the proliferation capacity of B-1a cells compared to B-2 cells in the peritoneal cavity. While we found no differences in BrdU incorporation between control and HuR KO peritoneal B-2 cells, a higher proportion of peritoneal HuR KO B-1a cells incorporated BrdU compared to control B-1a cells (***Fig. 3A***). Similarly, the percentage of splenic BrdU^+^ B-1a cells was higher in *Elavl1^fl/fl^ Mb1^Cre^* mice than in control mice (***Fig. 3B***). This, along with the fact that B-1 cell numbers are severely reduced in *Elavl1^fl/fl^ Mb1^Cre^* mice, suggests that HuR KO B-1a cells have a defect on cell maintenance rather than in cell proliferation.

**Figure 3.**
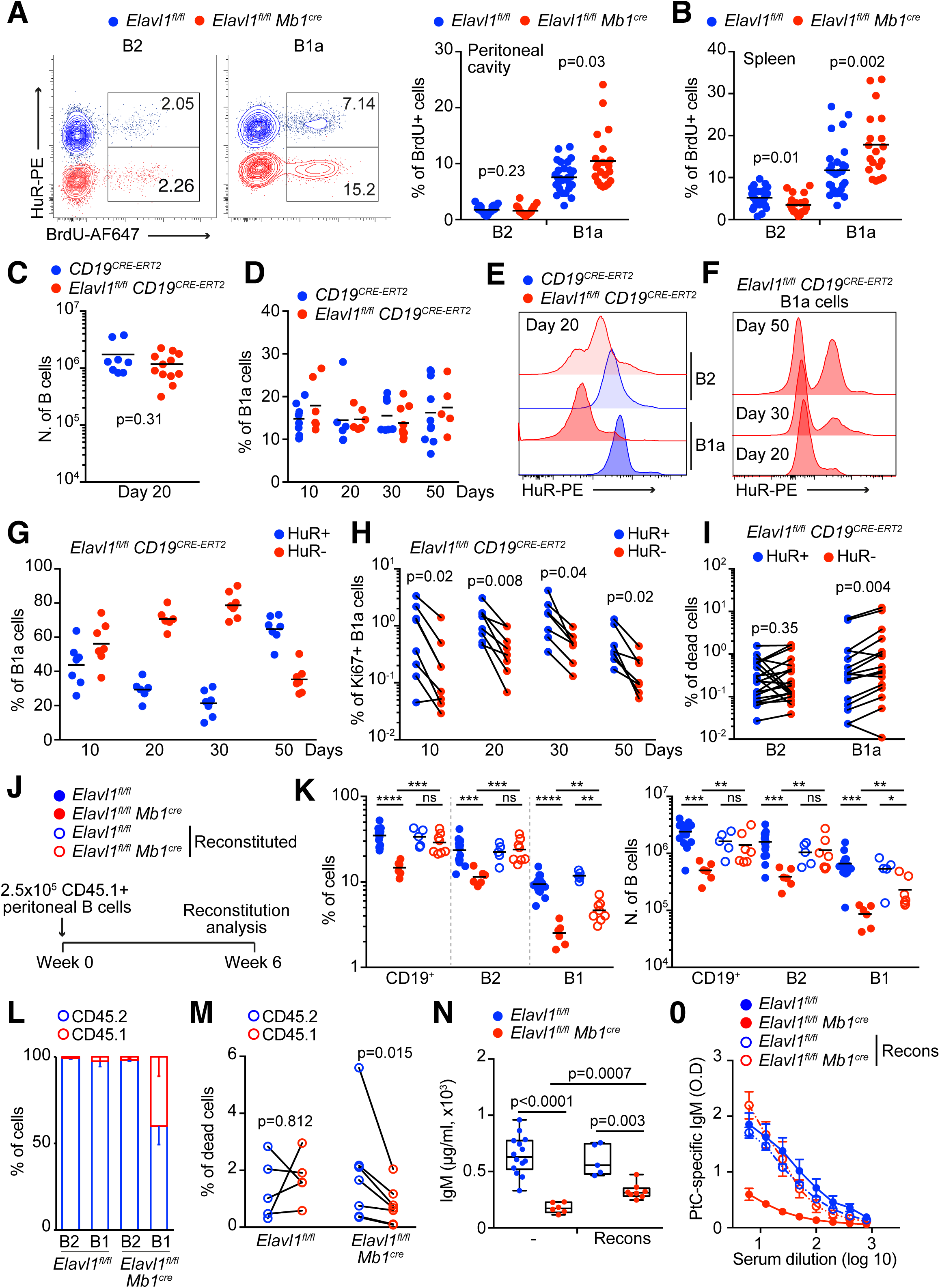
HuR is required for self-renewal and survival of innate B-1a cells. **A,** BrdU incorporation in peritoneal B-1a and B-2 cells. Left panels, representative contour plots showing BrdU staining in overlapped samples from control and *Elavl1^fl/fl^ Mb1^Cre^* mice. Right panel, quantitation of the percentage of BrdU^+^ B-2 and B-1a cells. Data pooled from three independent experiments performed each with at least 7 mice per genotype. **B,** Proportion of splenic BrdU^+^ B-2 and B-1a cells in mice shown in A. **C,** Number of peritoneal B cells in *CD19^CRE-ERT^*^2^ and *Elavl1^fl/fl^ CD19^CRE-ERT^*^2^ mice at day 20 after initiation of the tamoxifen treatment for 5 consecutive days. **D,** Percentage of B-1a cells in the peritoneal cavity of *CD19^CRE-ERT^*^2^ and *Elavl1^fl/fl^ CD19^CRE-ERT^*^2^ mice at the indicated days after treatment. **E,** Representative histogram showing HuR expression in peritoneal B-2 and B-1a cells from mice described in C. **F,** HuR expression in peritoneal B-1a cells from the *Elavl1^fl/fl^ CD19^CRE-ERT^*^2^ mice described in D. **G,** Proportion of HuR^+^ and HuR^-^ B-1a cells in the peritoneal cavity of *Elavl1^fl/fl^ CD19^CRE-ERT^*^2^ mice from D. **H,** Percentage of HuR^+^ Ki67^+^ and HuR^-^ Ki67^+^ B-1a cells in *Elavl1^fl/fl^ CD19^CRE-ERT^*^2^ mice. **I,** Quantitation of the percentage of B-2 and B-1a dead cells (FITC-VAD-FMK^+^ Zombie NIR Fixable Viability dye^+^) in the peritoneal cavity of *CD19^CRE-ERT^*^2^ and *Elavl1^fl/fl^ CD19^CRE-ERT^*^2^ mice treated with tamoxifen. Data shown in C to I were pooled from at two independent experiments performed each with at least 3 mice per genotype. **J,** Graphical summary of the experimental set up for peritoneal B cell reconstitution in *Elavl1^fl/fl^ Mb1^Cre^* mice. **K,** Percentage and number of total B, B-2 and B-1 cells in the peritoneal cavity of control and *Elavl1^fl/fl^ Mb1^Cre^* mice transferred or not with 250.000 CD45.1^+^ B cells from wild -type mice. Data representative from a representative experiment performed with at least 5 mice per group. **L,** Proportion of CD45.1^+^ and CD45.2^+^ cells identified within the peritoneal B-2 and B-1 cell subsets from mice in K. Data shown as mean ± SEM. **M,** Comparison of cell death between HuR-sufficient (CD45.1^+^) and HuR-deficient B-2 and B-1a cells 6 weeks after transplantation of peritoneal B cells in control and *Elavl1^fl/fl^ Mb1^Cre^* mice. **N,** IgM levels in the serum of mice shown in K. **O,** Serum dilution curves quantifying PtC-specific IgM antibodies in mice shown in K. ELISA data shown mean ± SEM. Two-tailed Mann-Whitney tests were performed for statistical analysis in A, B, C, K and N. Two-tailed paired Wilcoxon tests were used in H, I and M for statistical analysis.

To confirm the role of HuR in B-1a cell maintenance in a non-lymphopenic environment, we assessed the proliferation and death of B cells in *Elavl1^fl/fl^ CD19^CRE-ERT^*^2^ mice after treatment with tamoxifen for 5 consecutive days (***Suppl. Fig. 2A***). A first analysis confirmed that the number of total B cells and the percentage of B-1a cells remained constant in the peritoneal cavity of these mice independently of the elapsed time after tamoxifen treatment (***Fig. 3C*** and ***3D***). Further interrogation of the expression of HuR by flow cytometry showed a progressive loss of protein until day 30, time at which 80% of peritoneal cavity B-1a cells from tamoxifen-treated *Elavl1^fl/fl^ CD19^CRE-ERT^*^2^ mice lacked the expression of HuR (***Fig. 3E-3G***). By contrast, HuR deletion was less efficient in peritoneal B-2 cells and splenic B-1a cells, reaching up to 60% and 30% deletion efficiency respectively (***Suppl. Fig. 2B*** and ***2C***). Interestingly, the percentage of HuR^+^ B-1a cells remaining in the peritoneal cavity of *Elavl1^fl/fl^ CD19^CRE-ERT^*^2^ mice treated with tamoxifen jumped back from 20% at day 30 to 65% at day 50, suggesting that HuR-deficient B-1a cells could be outcompeted by those B-1a cells escaping tamoxifen-induced gene deletion (***Fig. 3F*** and ***3G***). Indeed, analysis of Ki67, a cellular marker expressed by cycling cells, revealed a higher proportion of HuR^+^ Ki67^+^ B-1a cells compared to HuR^-^ Ki67^+^ B-1a cells, both in the peritoneal cavity and the spleen (***Fig. 3H*** and ***Suppl. Fig. 2D***). Later measurement of cell death, by combining cell staining of active caspases with FITC-VAD-FMK with cell viability dyes, showed a higher proportion of B-1a dead cells in the absence of HuR (***Fig. 3I***). By contrast, the percentage of HuR^+^ and HuR^-^ B-2 dead cells remained similar. Taken together, these data show that HuR is essential for the homeostatic self-replenishment and survival of B-1a cells in mice.

Due to the self-replenishment capacity of B-1a cells, we hypothesized that the peritoneal B-cell compartment could be restored in *Elavl1^fl/fl^ Mb1^Cre^* mice by transferring a small fraction of HuR-sufficient B cells expressing the congenic marker CD45.1 for subsequent cell tracking (***Fig. 3J***). Indeed, quantitation of the percentage and number of B cells in *Elavl1^fl/fl^ Mb1^Cre^* mice reconstituted with solely 250.000 peritoneal B cells revealed a complete or partial restoration of the B-2 and B-1 cell compartment after 6 weeks of the cell transfer (***Fig. 3K***). While the restoration of the B-2 cell compartment was independent of exogenous cells, CD45.1 B-1 cells accounted for up to 40% of the total B-1 cell population in reconstituted *Elavl1^fl/fl^ Mb1^Cre^* mice (***Fig. 3L***). This percentage was below 5% in control mice similarly reconstituted with the same number of peritoneal CD45.1 B cells. Interestingly, cell survival analyses revealed that CD45.2 B-1 cells from the host *Elavl1^fl/fl^ Mb1^Cre^* mice died at a higher extend than transferred CD45.1 B-1 cells (***Fig. 3M***), reinforcing our previous observations on the need of HuR expression for B-1 cell survival. Quantitation of antibody levels in the serum of B-1 cell reconstituted *Elavl1^fl/fl^ Mb1^Cre^* mice also showed a partial restoration in total IgM production (***Fig. 3N***) and a complete rescue in the production of antibodies against PtC (***Fig. 3O***). In summary, we demonstrate that the B-1 cell compartment can be partially restored in *Elavl1^fl/fl^ Mb1^Cre^* mice by transferring HuR-sufficient B-1 cells holding a superior survival capacity to outcompete HuR KO B-1 cells.

### HuR shapes the transcriptome of innate B-1a cells

To gather insight into the molecular and cellular mechanisms regulated by HuR in innate B-1a cells, we designed a combined approach to investigate changes in the B cell transcriptome at the quantitative and qualitative level, and to identify the direct RNA targets of HuR in peritoneal cavity B cells using individual nucleotide cross-linking immunoprecipitation (iCLIP) (***Suppl. Fig. 3A***). Principal component analyses revealed that, while B-1a and B-2 cells from *Elavl1^fl/fl^ Mb1^Cre^* mice remained highly distinctive populations, the transcriptional differences between control and HuR KO B-1a cells were higher than those found when comparing control and HuR KO B-2 cells (***Fig. 4A***). Indeed, we identified three times the number of differentially expressed (DE) genes in HuR KO B-1a cells (1790 DE genes) than in HuR KO B-2 cells (672 DE genes), when compared to their control cell counterparts (***Fig. 4B*** and ***Suppl. Table 1***). More than 50% of DE genes in HuR KO B-2 cells were also found altered in HuR KO B-1a cells (341 genes). However, these genes represented only 20% of the DE genes found in HuR KO B-1a cells. Amongst the genes which expression was altered only in HuR KO B-1a cells, we found 52 *Ighv* and 32 *Igkv* genes representing 42% and 27% of the total of these genes (***Fig. 4C*** and ***Suppl. Table 1***). We could identify *Ig gene* pairings like *Ighv12-3/Igkv4-91* and *Ighv11-2/Igkv14-126* as being drastically reduced in HuR KO B-1a cells. This was in line with the impaired expansion of common B-1 cell clonotypes recognising PtC and other self-antigens in *Elavl1^fl/fl^ Mb1^Cre^* mice.

**Figure 4.**
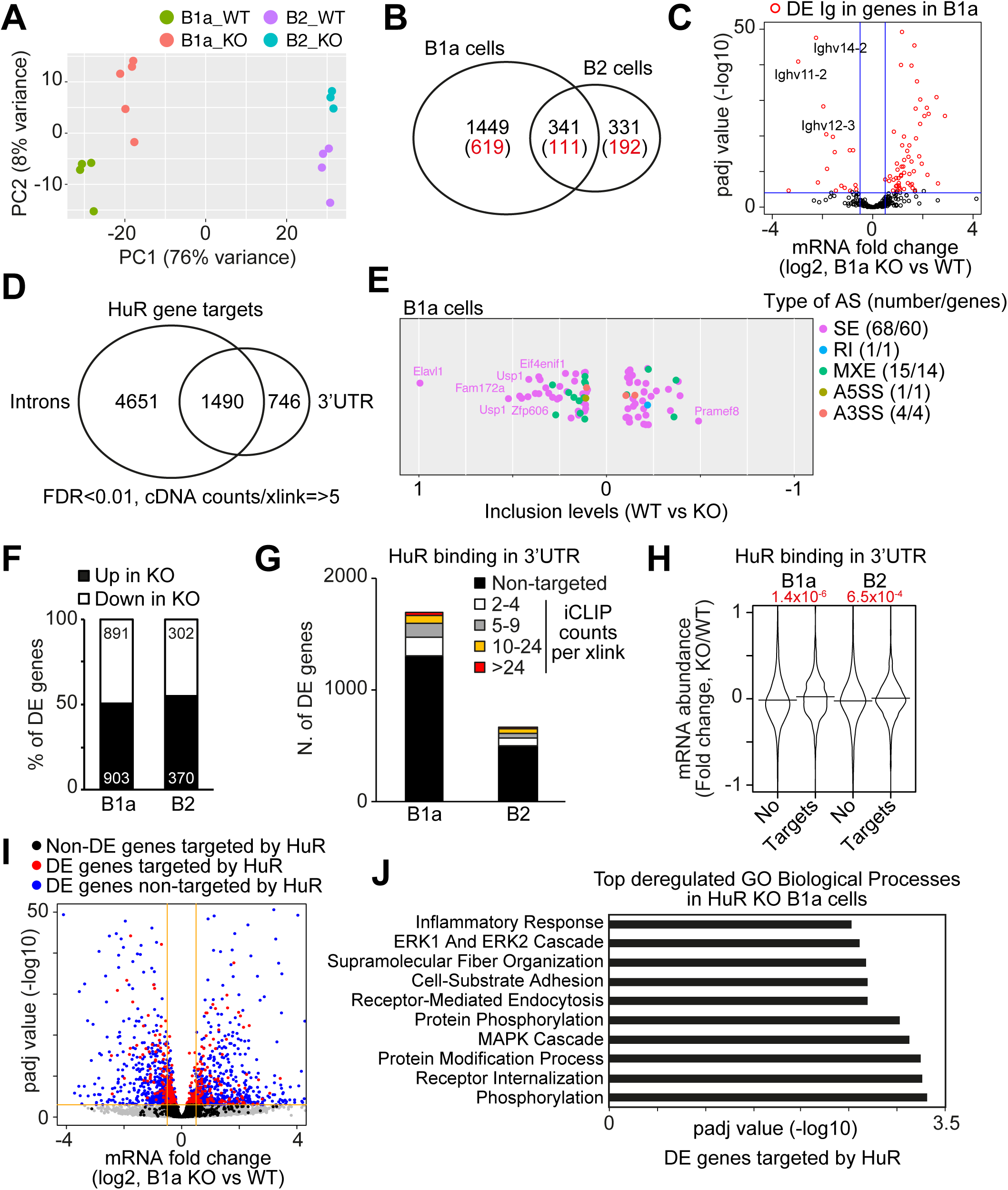
HuR shapes the transcriptome of innate B-1a cells. **A,** Principal component analysis comparing the transcriptome of B-1a and B-2 cells from control and *Elavl1^fl/fl^ Mb1^Cre^* mice. **B,** Venn-diagram showing the number of differentially expressed (DE) genes in peritoneal HuR KO B-1a and B-2 cells when compared to their control counterparts. In black, DE genes with FDR<0.05. In red, DE genes with FDR<0.05 and an absolute fold change higher than 2. **C,** Volcano plot showing the fold change in abundance of *Ighv* mRNAs in HuR KO B-1a cells compared to control B-1a cells. In red, *Ighv* mRNA changes with FDR<0.05 and an absolute fold change higher than 1.5. **D,** Venn-diagram of HuR target genes in which HuR crosslinks are annotated within introns or 3’UTRs. HuR crosslinks are defined by at least 5 unique cDNA counts and FDR<0.01. **E,** Alternative splicing events in HuR KO B-1a cells uncovered by rMATS. Colour code indicates the different types of splicing events (SE: spliced exon, RI: retained intron, MXE: mutually exclusive exon, A5SS: alternative 5’ spliced site and A3SS: alternative 3’ spliced site). Left number: number of events. Right number: number of genes affected. **F,** Proportion of DE genes that are either up- or down-regulated in HuR KO B-1a and B-2 cells. **G,** Number of DE genes that are targets of HuR. **H,** Global analysis of mRNA abundance for genes targeted or not by HuR. Kolmogorov– Smirnov tests are performed to assess changes in data distribution between HuR-targets and non-targets in B-1a or B-2 cells. **I,** Volcano plot showing changes in mRNA abundance of HuR targets in HuR KO B-1a cells. **J,** Top-10 pathways containing the genes targeted by HuR and deregulated in HuR KO B-1a cells. Pathway enrichment analysis was performed with Enrichr^79^. Pathway database, GO Biological Processes.

HuR can shape the cell transcriptome and proteome by controlling mRNA splicing, stability and/or translation. Using iCLIP, we found that HuR binds preferentially to introns and 3’ untranslated regions (3’UTRs) in peritoneal cavity B cells (***Suppl. Fig. 3B***). The vast majority of HuR:RNA crosslinks (59%) were annotated within the intronic regions of 6141 genes (***Fig. 4D*** and ***Suppl. Fig. 3C***) (***Suppl. Table 2***). However, HuR crosslink density was highest in 3’UTRs (***Suppl. Fig. 3D***), with 8% of total HuR:RNA crosslinks mapped to the 3’UTRs of 2236 genes (***Fig. 4D*** and ***Suppl. Fig. 3C***). To uncover a putative role of HuR as splicing regulator, we quantified changes in mRNA splicing between control and HuR KO B-1a cells using rMATS. We could annotate only 89 alternative splicing events associated to 80 genes in the absence of HuR (***Fig. 4E***). Among these few events, we could identify 100% exclusion of *Elavl1* exon 2, which is flanked by loxP sites and deleted upon Cre-recombination in *Elavl1^fl/fl^ Mb1^Cre^* mice. This highlights the capacity of rMATS to detect changes in mRNA splicing and allows us to conclude that HuR is not a major splicing regulator in resting innate B-1a cells. Next, we assessed whether HuR could act as a global modulator of mRNA stability. HuR KO B-1a and B-2 cells showed a similar proportion between DE genes up- and down-regulated in the absence of HuR (***Fig. 4F***). Integration of HuR:RNA interactome data revealed that only 25% of DE genes in both HuR KO B-1a and B-2 cells were directly targeted by HuR (***Fig. 4G***). Further comparison of global changes in mRNA abundance showed that HuR targets were mildly, but significantly, increased in HuR KO B-1a and B-2 cells (***Fig. 4H***). Thus, we conclude that HuR is essential for the correct expression of thousands of genes in innate B-1a cells, but it does not act as a global regulator of mRNA splicing or stability.

### HuR controls BCR expression and BCR-dependent tonic signalling

Genes targeted by HuR showed milder differences in gene expression than those non-targeted by HuR (***Fig. 4I*** and ***Suppl. Fig. 3D***). This suggested that HuR could be controlling the expression of few master gene regulators which, later, contribute to the ample remodelling of the B-1a cell transcriptome. Indeed, gene enrichment analyses revealed that phosphorylation, receptor internalization and the ERK/MAPK cascades were the most altered pathways in HuR KO B-1a cells driven by HuR target genes (***Fig. 4J*** and ***Suppl. Table 3***).

ERK/MAPK and PI3K/AKT pathways are major signalling hubs activated during tonic activation of the B-cell receptor (BCR) and, if altered, they compromise the homeostatic maintenance and self-replenishment of innate B-1a cells ^44, 45^. Thus, we assessed in detail if BCR expression and/or BCR-dependent signalling was altered in HuR KO B-1a cells. B-1a cells expressed twice the amount of IgM than B-2 cells in control mice (***Fig. 5A***). However, both total and cell surface levels of IgM were significantly decreased in HuR KO B-1a cells (***Fig. 5A***). This defect was restricted to B-1a cells as it could not be observed in B-2 cells from *Elavl1^fl/fl^ Mb1^Cre^* mice. Next, we assessed the functional relevance of decreased IgM surface expression in HuR KO B-1a cells and measured by phospho-specific flow cytometry the phosphorylation of SYK, PLCγ2 and AKT, three essential signal transducers activated downstream the BCR. While basal levels of phospho-SYK, phospho-PLCγ2 and phospho-AKT remained similar in non-activated control and HuR KO B-1a cells, phosphorylation of these three signal transducers after BCR stimulation was significantly reduced in the absence of HuR (***Fig. 5B***, ***5C and 5D***). By contrast, BCR-dependent phosphorylation of PLCγ2 and SYK was normal in control and HuR KO B-2 cells, which showed only mild-alterations in the activation of the PI3K/AKT pathway (***Suppl. Fig. 4A***). Defective BCR-mediated signalling was confirmed in innate B-1a cells from the spleen of *Elavl1^fl/fl^ Mb1^Cre^* mice (***Suppl. Fig. 4B***). Importantly, expression of *Plcg2* and *Syk* remained unchanged in HuR KO B-1a cells (***Suppl. Table 1***); and stimulation with pervanadate, a powerful inhibitor of protein tyrosine phosphatases (PTP) that act downstream the BCR to limit exacerbated B cell activation, revealed similar levels of pPLCγ2 and pSYK in control and HuR KO B-1a cells (***Suppl. Fig. 4C***). Taken together, our data highlight that HuR controls high surface expression of IgM in B-1a cells and, consequently, downstream BCR-dependent homeostatic signalling.

**Figure 5.**
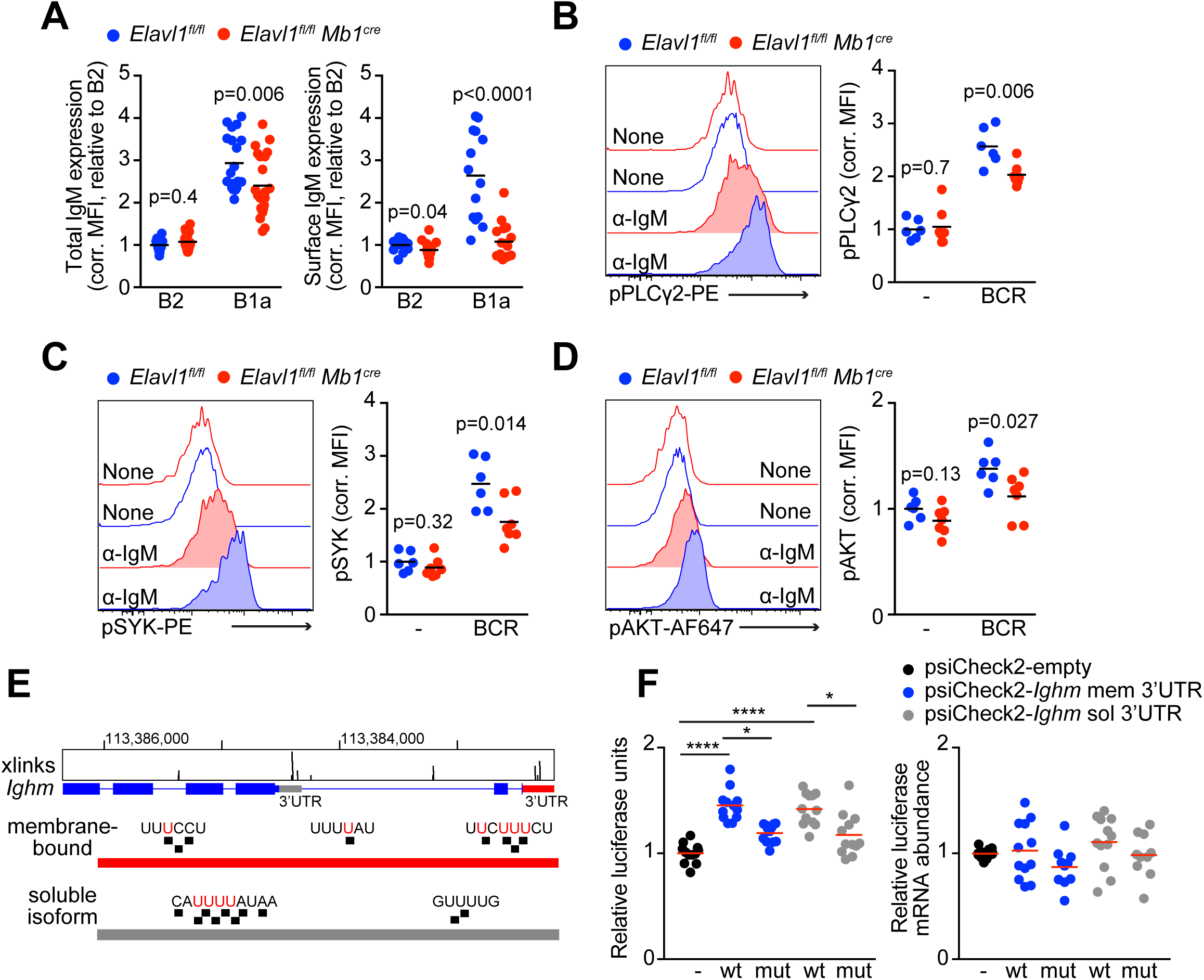
HuR controls BCR expression and homeostatic signalling. **A,** Total and surface IgM levels in peritoneal B-1a and B 2 cells from control and *Elavl1^fl/fl^ Mb1^Cre^* mice measured by flow cytometry. Data pooled from four independent experiments performed with at least 3 mice per genotype. **B, C, D,** Phospho-flow cytometry analysis of pPLCγ2 (B), pSYK (C) and pAKT (D) in peritoneal B-1a cells stimulated in-vitro with an F(ab’)2-anti-IgM antibody for 5 min. Left panels, representative histograms showing the expression of phospho-proteins in non-treated and treated B-1a cells from control and *Elavl1^fl/fl^ Mb1^Cre^* mice. Right panels, quantitation of phospho-proteins normalised by the levels in control B-1a cells non-treated. Data pooled from two independent experiments performed with at least 3 mice per genotype. **E,** Genomic view of HuR crosslinks annotated to the 3’UTRs of membrane-bound and soluble-bound *Ighm* mRNA transcripts. Bases mutated in 3’UTR luciferase reporter assays are indicated in red. **F,** Quantitation of *Renilla* luciferase translation (left panel) and stability (right panel) under the control of the 3’UTRs of the soluble- or membrane-bound *Ighm* transcript isoforms. Wild-type (wt) or HuR-binding site mutated (mut) 3’UTRs are compared. *Renilla* luciferase units were normalized by translation of *Firefly* luciferase, that acted as an internal control, and were relative quantified to an empty psiCheck2 reporter vector lacking a 3’UTR. Data pooled from three independent experiments performed each of them by triplicate. *Renilla luciferase* mRNA abundance was measured by RT-qPCR and normalised by the expression of 18S rRNA and firefly luciferase to correct for differences in starting material and transfection efficiencies, respectively. Two-tailed Mann-Whitney tests were performed for statistical analysis in A, B, C and D. One-way ANOVA and Tukey’s multiple comparison tests were performed in F (* p<0.05, **** p<0.0001).

To gather insight into the molecular mechanism by which HuR controlled IgM expression, we interrogated the binding properties of HuR with *Ighm* mRNAs in peritoneal B cells. HuR:RNA interactomics data showed that HuR physically interacted with U-rich elements present in the 3’UTRs of *Ighm* transcripts encoding both soluble- (*Ighm-μs*) and membrane-IgM (*Ighm-μm*) (***Fig. 5E*** and ***Suppl. Table 2***). Therefore, we constructed functional assays in which the 3’UTR of these two transcript isoforms were cloned downstream a renilla luciferase reporter. 3’UTRs from *Ighm-μs* and *Ighm-μm* transcripts promoted mRNA translation without altering mRNA abundance. Mutation of the HuR-bound uridines identified with iCLIP significantly reduced the translation, but not the mRNA stability, of the luciferase reporter under the regulation of the *Ighm* 3’UTRs (***Fig. 5F***). This leads us to conclude that physical interaction of HuR to *Ighm* mRNA transcripts promotes their translation into protein in innate B-1 cells.

### HuR supports BAFF/APRIL-dependent B cell survival

RBP-dependent control of functionally related mRNAs allows the regulation of biological functions in time and magnitude ^29^. Along with tonic BCR signalling, co-activation of the surface receptors TACI and BAFFR contributes to the survival of B cells ^46^. HuR:RNA interactomics in peritoneal B cells identified *Tnfrsf13b* and *Tnfrsf13c*, the mRNAs encoding for TACI and BAFFR respectively, as targets of HuR (***Fig. 6A***). In line with previous reports, TACI and BAFFR were highly expressed in B-1a cells compared to B-2 cells (***Fig. 6B*** and ***6C***). Importantly, total and surface expression of these receptors were significantly reduced in B-1a and B-2 cells from *Elavl1^fl/fl^ Mb1^Cre^* mice when compared to control littermates. *Tnfrsf13b* and *Tnfrsf13c* mRNA abundance was also reduced in HuR KO B-1a cells (***Suppl. Table 1***), indicating that HuR is likely implicated in the stabilization and/or translation of *Tnfrsf13b* and *Tnfrsf13c* mRNAs.

**Figure 6.**
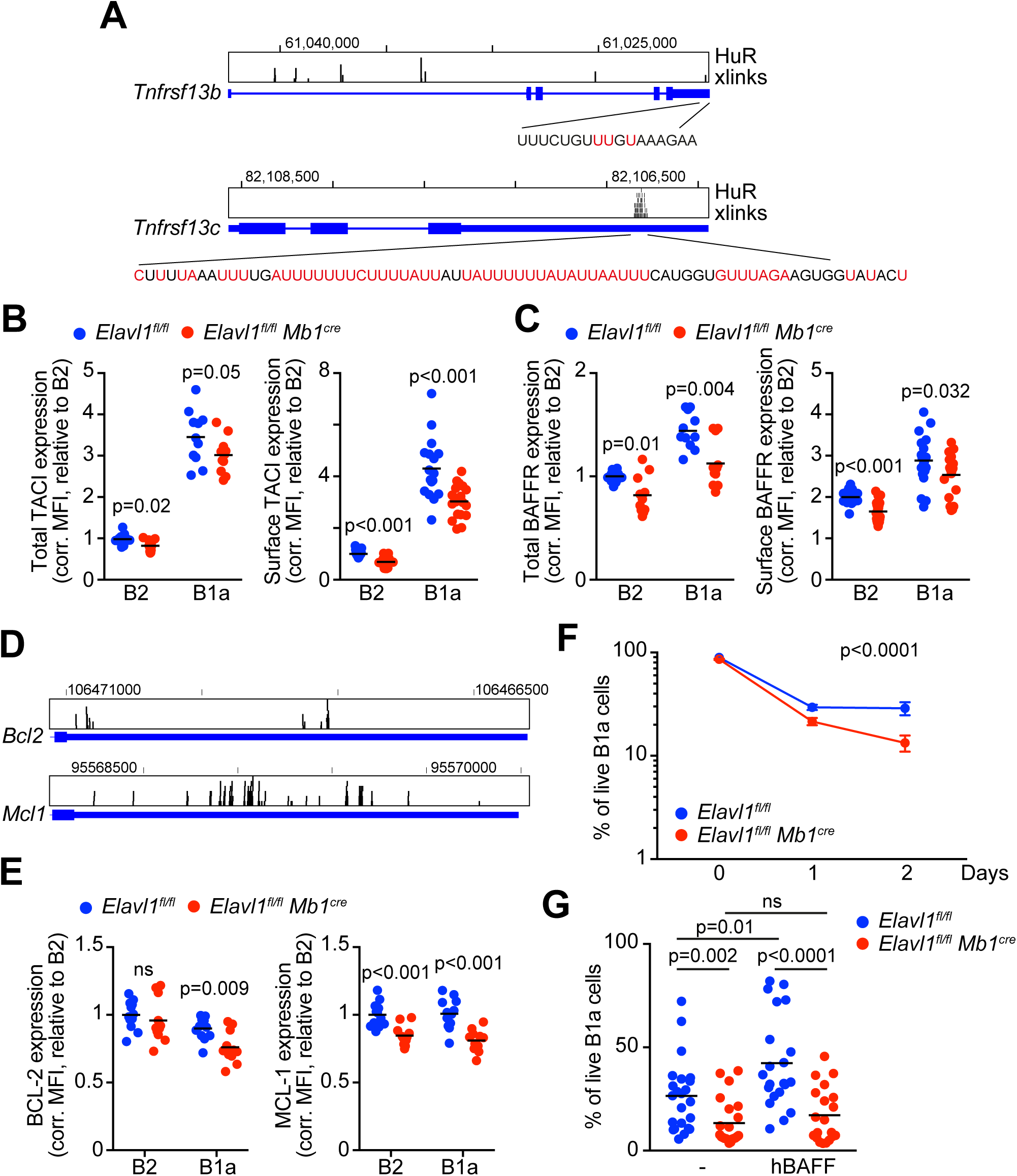
HuR enables B-1a cell survival. **A,** Genomic view of HuR binding to the 3’UTRs of *Tnfrsf13b* and *Tnfrsf13c* in peritoneal B cells. Unique crosslink sites are highlighted in red. **B, C,** Quantitation by flow cytometry of the total and surface levels of TACI and BAFFR in peritoneal B-1a and B-2 cells from control and *Elavl1^fl/fl^ Mb1^Cre^*mice. Data pooled from three independent experiments performed with at least 3 mice per genotype. **D,** Visualization of HuR binding to the 3’UTRs of *Bcl2* and *Mcl1* in peritoneal B cells. **E,** Quantitation of BCL2 and MCL1 protein expression in peritoneal B-2 and B-1a cells. Data pooled from two independent experiments performed with at least 4 mice per genotype. **F,** Assessment of peritoneal B-1a cell viability after *in-vitro* culture for 1 or 2 days. Viability was measured by flow cytometry after cell staining with Dapi. **G,** Proportion of live B-1a cells after in-vitro culture for 2 days in the presence of hBAFF. Data shown in D and E were pooled from four independent experiments in which at least 3 mice per genotype were cultured independently. Two-tailed Mann-Whitney tests were performed for statistical analysis in B, C, E and G. Two-way ANOVA was performed in F.

BAFF/APRIL signalling contributes to B-cell maintenance by promoting the expression of the pro-survival molecules BCL-2 and MCL-1 while restraining apoptotic drivers such as BIM ^47,48^. To uncover the functional consequences of HuR deficiency and reduced levels of TACI and BAFFR in the expression of BCL-2 and MCL-1, we first confirmed previous observations describing the specific interaction of HuR with AU-rich elements present in the 3’UTR of *Bcl2* (***Fig. 6D***) ^49^. Additionally, HuR:RNA interactomics analyses revealed an extensive association of HuR along the 3’UTR of *Mcl1* (***Fig. 6D***) suggesting the existence of an HuR-dependent regulatory mechanism controlling the expression of both BCL-2 and MCL-1. *Bcl-2* and *Mcl-1* mRNA abundance remained unchanged in peritoneal HuR KO B cells compared to control B cell counterparts (***Suppl. Table 1***). However, we found protein levels of BCL-2 and MCL-1 significantly diminished in the absence of HuR (***Fig. 6E***) and a concomitant reduction in HuR KO B-1a cell viability when compared to control B-1a cells (***Fig. 6F***). Addition of soluble BAFF into the culture media increased the survival capacity of control B-1a cells, but failed to rescue HuR KO B-1a cell viability (***Fig. 6G***). Taken together, our data demonstrate that HuR is essential for BAFF/APRIL-dependent B-1a cell survival by fine-tuning the expression of TACI, BAFFR, BCL-2 and MCL-1.

### Expansion of self-antigen specific B cells is impaired in the absence of HuR

Recognition of self-antigen phospholipids by innate B-1 cells allows direct removal of apoptotic cells and the production of autoantibodies for enhanced recognition and phagocytosis of these apoptotic bodies by other immune cells ^50^. Optimal expression of BAFFR and, specially, TACI is required for similar T-independent type II antibody responses ^51, 52^. Thus, to explore the functional consequences of HuR deletion in innate B cell-driven immunity, we immunized 8-12-week-old control and *Elavl1^fl/fl^ Mb1^Cre^* mice with thymocyte-derived apoptotic bodies and assessed the expansion of total and PtC-specific B-1 cells (***Fig. 7A*** and ***7B***). In general, we recorded a tendency for innate B-1 cells to accumulate in the peritoneal cavity of control mice after immunization (***Fig. 7C***). By contrast, the percentage of B-1 cells in *Elavl1^fl/fl^ Mb1^Cre^* mice remained 2.5-fold lower than in control mice independently on whether there were immunised or not (***Fig. 7C*** and ***Suppl. Fig 5A***). While PtC-specific B cells almost double in control mice upon immunization, this population remained highly reduced in *Elavl1^fl/fl^ Mb1^Cre^* mice (***Fig. 7B*** and ***Fig. 7D***). Over 90% of PtC-specific B cell clones in the peritoneal cavity originated from B-1 cells (***Suppl. Fig 5B***), and constituted up to 50% of the B-1 cell compartment in immunized control mice (***Suppl. Fig 5C***). This proportion was significantly lower in *Elavl1^fl/fl^ Mb1^Cre^* mice and did not change upon mouse immunization with apoptotic bodies. As a consequence, serum levels of IgM remained barely detectable in *Elavl1^fl/fl^ Mb1^Cre^* mice and PtC-specific IgM did not increase upon immunization as in control mice (***Fig. 7E*** and ***Fig. 7F***). In summary our data suggest that, upon immunization with apoptotic bodies, HuR is required for either the expansion or maintenance of PtC-specific B-1 cells and, in its absence, *Elavl1^fl/fl^ Mb1^Cre^* mice fail to produce significant amounts of autoantibodies.

**Figure 7.**
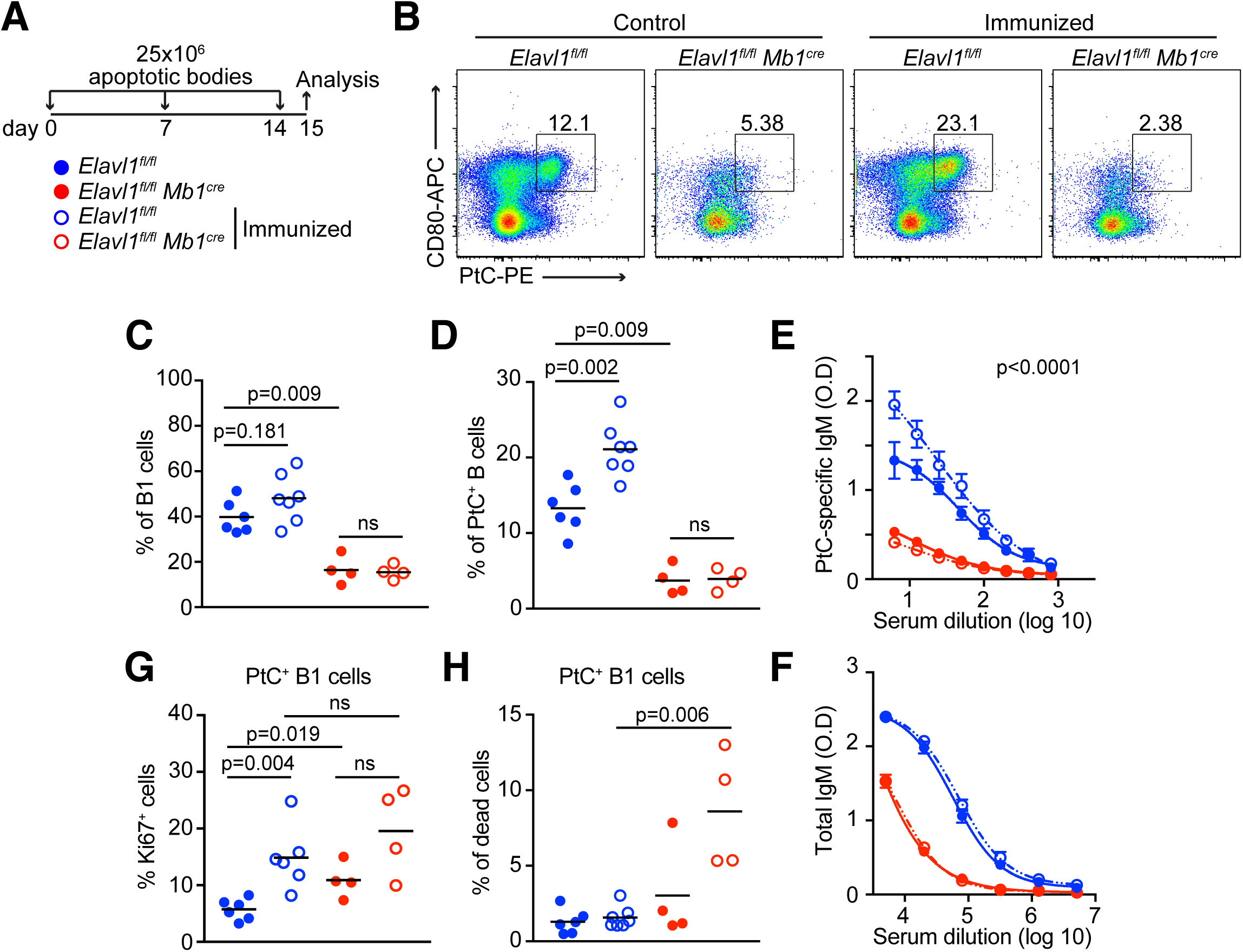
HuR is required for the expansion of self-antigen specific B-1a cells. **A,** Summary of B-1 cell expansion analysis upon immunization with thymic-derived apoptotic bodies. **B,** Representative pseudo-colour dot plots showing a gating strategy to detect PtC^+^ B-1 cells in immunized control and *Elavl1^fl/fl^ Mb1^Cre^* mice (gated on CD19^+^ cells). **C, D,** Proportion of total peritoneal B-1 cells (C) and PtC-specific B cells (D) in control and *Elavl1^fl/fl^ Mb1^Cre^* mice immunized or not with apoptotic bodies. Data representative from one of the experiments performed with a minimum of 4 mice per genotype. **E, F,** Serum dilution curves to quantify PtC-specific or total IgM antibodies in mice from C. Data shown mean ± SEM. Sum of squares F test was performed to assess changes in the curve data distribution between unimmunized and immunized control mice. **G,** Quantitation of the percentage of Ki67^+^ PtC-specific B-1 cells in mice from C. **H,** Analysis of PtC^+^ B-1 cell dead in mice from C measured by cell labelling with a Zombie NIR viability die. Two-tailed Mann-Whitney tests were performed in C, D, G and H for statistical analysis of the data.

Analysis of Ki67^+^ PtC-specific B-1 cells in control mice revealed a significant increase in the proportion of cycling cells upon immunization, as expected from B cells getting activated by cognate antigens (***Fig. 7G***). In line with our previous observations, the percentage of PtC-specific cycling B-1 cells was increased in unimmunised *Elavl1^fl/fl^ Mb1^Cre^* mice compared to control mice and it did not augment further upon immunization. Finally, quantitation of cell death using viability dies revealed a profound increase in the percentage of PtC-specific B-1 dead cells in immunized *Elavl1^fl/fl^ Mb1^Cre^* mice compared to control mice (***Fig. 7H***). Altogether, these data lead us to conclude that PtC-specific HuR KO B-1 cells can get activated and enter the cell cycle normally, although they fail to expand in response to self-antigens due to increased cell death.

## DISCUSSION

B-1 cells constitute a critical immunoregulatory layer for self-defence which origin and maintenance are far from being fully understood. Here we provide insight into how post-transcriptional gene regulation by HuR shapes the innate B-1 cell compartment by promoting BCR-dependent survival and expansion. Our data demonstrate that HuR is dispensable during early B-1 cell development. However, absence of HuR leads to a drastic reduction in the number of mature B-1 cells and the production of natural autoantibodies. Mechanistically, HuR contributes to the homeostatic maintenance and self-replenishment of B-1 cells by securing the expression of the receptors BCR, TACI and BAFFR, and their downstream signalling.

Innate B-1 cells produce up to 90% of low-affinity and polyreactive serum IgM antibodies ^53, 54^, with a repertoire skewed to the recognition of invariant constituents from microbial and apoptotic cell membranes (e.g. PC and PtC). Interaction of such antibodies with these and other autoantigens removes noxious molecules prior they trigger an exacerbated immune response and irreversible tissue damage. *Elavl1^fl/fl^ Mb1^Cre^* mice produce only around 10% of the normal amount of total, and PC- and PtC-specific IgM antibodies found in the serum of adult mice. This decrease could be circumscribed not only to the large reduction in the numbers of B-1 cells found in *Elavl1^fl/fl^ Mb1^Cre^* mice but, more importantly, with the expression of a neonate-like BCR repertoire in these HuR KO B-1 cells. It has been proposed that, as the mouse aged, recurrent detection of self-antigens from dead cells couples mature B-1 cell replenishment with restrictive clonal selection ^3, 7, 8^. Indeed, our data confirm previous observations that associate the emergence of common BCR clonotypes against PC and PtC with age ^9^. This selection process can be speeded up in control mice by repetitive immunization with apoptotic cell bodies. However, *Elavl1^fl/fl^ Mb1^Cre^* mice fail to expand their PtC-specific B-1 cell population both during aging and after immunization. This is despite the fact that, in these mice, lymphopenia in the peritoneal cavity increases cycling of HuR KO B-1 cells in order to fill the innate B cell pool. By contrast, in a non-lymphopenic environment, HuR KO B-1 cells show a profound competitive disadvantage in the presence of HuR-sufficient B-1 cells. These wild-type cells are capable to fully rescue serum levels of PtC-specific antibodies in just 6 weeks after adoptive transfer into *Elavl1^fl/fl^ Mb1^Cre^* mice. Our data demonstrate that HuR promotes B-1 cell proliferation and survival, in line with previous observations in other cell types ^55, 56^. Mechanistically, HuR critically contributes to the expression of the cell surface receptors BCR, TACI and BAFFR, which are all required for downstream MAPK, PI3K and NFkB signalling activation and proliferation of B cells ^28, 46, 57^. Transcripts encoding these cell surface receptors along with *Bcl-2* and *Mcl-1* mRNAs constitute an HuR-dependent RNA operon indispensable for B-1 cell maintenance and self-replenishment.

Analysis of the transcriptional changes in B-2 and B-1a cells in the absence of HuR reveals the large implication of this RBP in the homeostatic activation of B-1a cells in the peritoneal cavity. We could identify multiple pathways related to receptor activation, MAPK signalling cascade and protein phosphorylation as being deregulated in HuR KO B-1a cells. Interestingly, the impact of HuR deletion is milder in B-2 cells, with none of the gene signatures indicated above being significantly deregulated in these quiescent cells. This is in line with our previous observations showing that HuR-dependent regulation increases upon *in-vitro* activation of follicular B cells and in germinal centre B cells ^40, 42^.

HuR binds to U- and AU-rich regulatory elements present in introns and 3’UTRs. However, rather than acting as a splicing modulator, HuR mainly controls mRNA translation in B-1a cells. The low number of B-1a cells recovered from *Elavl1^fl/fl^ Mb1^Cre^* mice impeded us to assess global translation using methodologies such as ribosome footprinting. Nonetheless, using iCLIP and 3’UTR luciferase reporter assays, we demonstrate that HuR binds to the 3’UTR of both *Ighm-μs* and *Ighm-μm* transcripts and promotes their translation into protein.

IgM (BCR) surface expression is decreased by almost 3-fold in HuR KO B-1a cells. This defect translates into a defective phosphorylation of downstream signal transducers PLCγ2, SYK and AKT, three key components of the BCR signalosome that link BCR-antigen engagement with activation of MAPK, PI3K and NF-kB signalling pathways ^58^. Importantly, B-cell treatment with pervanadate, a phosphatase inhibitor that blocks constitutive negative regulation of BCR signalling exerted by SHP-1/-2, SHIP-1/-2 and PTEN ^58^ ^59^, reveals that global phosphorylation capacity of PLCγ2 and SYK in B-1a cells remains unaffected in the absence of HuR. Similarly, the mRNA abundance of *Plcg2* and *Syk* is comparable between control and HuR KO B-1a cells. Therefore, we conclude that altered BCR-dependent signalling in HuR KO B-1a cells is mostly a reflection of a defective *Ighm* mRNA translation rather than global alterations in downstream signal transducers.

Instructive BCR-signalling is required for B-1 cell development and selection but, in the absence of co-stimulatory signals, leads into cell death. Integration of multiple signalling pathways is required in B cells, with the cytokines BAFF and APRIL providing a must needed signal for B cell metabolic fitness and survival ^46, 60, 61, 62, 63^. These cytokines are recognised by the receptor BAFFR, TACI and BCMA, with the latest being absent in B-1a cells. A previous study suggested that HuR can modulate BAFFR subcellular location without altering its protein abundance ^64^. However, our data show that HuR is required for both mRNA and protein expression of BAFFR and TACI in peritoneal B cells. Further analyses, showing a reduction in total and cell surface expression of these receptors, suggest that HuR controls their mRNA stability and/or translation rather than alternative 3’UTR usage and membrane protein location in peritoneal B cells.

BAFF-mediated modulation of pro- and anti-apoptotic members of the BCL2-family of proteins sets the capacity of B cells to survive in the periphery. The absence of MCL-1 or BCL-2 in B cells, or just deletion of the AU-rich regulatory elements in the 3’UTR that controls *Bcl2* mRNA stability and translation, leads to a reduction in the number of these cells in peripheral organs ^49, 65, 66^. MCL1 and BCL2 proteins, but not their mRNA transcripts, are significantly decreased in HuR KO B-1a cells. This confirms previous observations in cell lines showing that HuR-binding to *Mcl1* and *Bcl2* 3’UTRs fine-tunes the expression of these anti-apoptotic molecules ^67, 68^. By controlling the expression of BCR and survival receptors, and anti-apoptotic molecules, HuR orchestrates an integrative program for long-lasting maintenance and self-renewal of peritoneal B-1 cells.

Coordinated regulation of functionally related mRNA operons by RBPs is a powerful molecular mechanism that defines the fate of cells from development to function ^69^. Several operons can co-exist in the same cell ^29^. This is exemplified in progenitor B cells by the fact that while TIA1 and TIAL1 control the splicing of multiple mRNAs of the DNA damage repair machinery ^32^, the RBPs ZFP36L1 and ZFP36L2 modulates the stability of an mRNA operon controlling cell quiescence and proliferation ^37^. Similarly, RBPs such as HuR can either modulate the same mRNA operon in different cell types or different ones depending on the cell type or activation status. This study provides another piece of evidence about the importance of coordinated post-transcriptional regulation in the establishment and function of B cells. In the future, uncovering how transcriptional and post-transcriptional gene programs are intertwined in B cells should bring light into the complex mechanisms that build our immune system against external and self-antigens.

## Acknowledgments

We thank to all personnel from the animal facility of Toulouse-CREFRE, and from the flow cytometry (F. L’Faqihi, V. Duplan-Eche and AL. Iscache), transcriptomics (A. Chaubet) and bioinformatics technical platforms of INFINITY. This project was supported by the ATIP-Avenir, Plan Cancer program (C18003BS), the French National Research Agency, ANR (ANR-20-CE15-0007, ANR-22-CE15-0013 and ANR-24-CE15-5291), FRM Foundation (EQU202303016269, FDT202304016365), the ARSEP foundation (R19201BB and 1299) and Boehringer Ingelheim (BIF) Fonds.

## CRediT author contributions

Dunja Capitan-Sobrino: Conceptualisation, Investigation, Formal analysis, Funding acquisition, Writing - review and editing. Mailys Mouysset: Investigation, Formal analysis, Writing - review and editing. Orlane Maloudi: Investigation, Formal analysis. Yann Aubert: Formal analysis, Writing - review and editing. Ines C. Osma-Garcia: Investigation, Formal analysis, Funding acquisition, Writing - review and editing. Maia Nestor-Martin: Investigation, Formal analysis. Trang-My M. Nguyen: Investigation, Formal analysis. Greta Dunga: Investigation, Formal analysis. Manuel D. Diaz-Muñoz: Conceptualisation, Investigation, Formal analysis, Resources, Supervision, Funding acquisition, Writing — original draft, review and editing, Project administration.

## Ethics declaration

The authors declare no competing interests.

## METHODS

### Animal models

*Elavl1^tm^*^1d^*^kon^*(*Elavl1^fl/fl^*) mice ^70^ were crossed with *CD79a^tm^*>^1^(cre)*^Reth^*(*Mb1^Cre^*) mice ^43^ or *B6.129P2-Cd19^tm^*>^1^(cre/ERT2)*^Rsky^/J* (from The Jackson Labs, *CD19^ERT^*^2^*^-Cre^*) for conditional gene deletion in newly developed B cells or after treatment with tamoxifen. C57BL/6-Ly5.1 (CD45.1^+^) mice were kindly provided by Dr. R. Liblau, N. Blanchard and Dr. F. Masson (INFINITY, Toulouse, France). All mice were maintained in a C57BL/6 genetic background. Both males and females were used indistinctly with the exception of analyses assessing self-reactive B-1 cells producing self-antibodies in which females were used. Randomization was set in all our studies by housing both Cre+ and Cre-control littermates in the same cage. Experimental ‘blinding’ was established in all experiments performed with embryos and pups with an age of 2-weeks or less as indicated below. For animals older than 2 weeks, genotyping was performed early on and experimental blinding was not established. No primary pathogens or additional agents listed in the FELASA recommendations were detected during health monitoring. Animal housing was kept at a constant temperature of ∼19-21C and a relative humidity of 52% at the Inserm service unit UMS006 (CREFRE, Toulouse, France). 3-5 animals were housed in ventilated cages with lighting provided on 12-hour light and 12-hour dark cycles. Mice were fed ad libitum and provided with environmental enrichment throughout their lifespan. All experimental procedures were approved by the local ethical committee of INFINITY and by the French Ministry of Education, Research and Innovation.

### Animal procedures

B cell populations were analysed from the peritoneal cavity, spleen and liver. Briefly, after CO2 euthanasia and terminal bleeding by cardiac puncture, peritoneal cavity cells were isolated by gentle washing of the peritoneal cavity with 10-15 ml of ice-cold PBS with 2% FCS. For spleen and liver, single cell suspensions were obtaining after tissue smashing through a 40 µm cell strainer. Any red blood cells were lysed using an ACK lysis buffer prior downstream applications. In some cases, mice were fed with BrdU (1 mg/ml; Sigma-Aldrich) / 1% sucrose or glucose in the drinking water ad libitum for 5 consecutive days prior euthanasia. Inducible deletion of HuR in *Elavl1^fl/fl^ CD19^ERT^*^2^*^-Cre^*and *CD19^ERT^*^2^*^-Cre^*mice was achieved after administering by oral gavage during 5 consecutive days 200 mg. of tamoxifen/kg body weight resuspended in sunflower oil and 10% ethanol. HuR deletion was assessed by flow cytometry at all experimental endpoints.

For B-1 cell embryonic development analysis, breeding females were monitored daily searching for the presence of a vaginal plug marking mating day. Females were euthanised 17.5 days later to collect E17.5 embryos. Both pregnant females and embryos were culled using C0_2_ or decapitation after anesthesia. Breeding strategy allowed generation of control and HuR cKO mice in a 50% mendelian-rate. The genotype of the embryos and new born mice (up to 2-weeks of age) was determined at the experimental endpoint by measuring HuR expression by flow cytometry.

Homeostatic replenishment of the B-1 cell compartment in *Elavl1^fl/fl^ Mb1^Cre^* mice was assessed after i.p. transfer of 2.5×10^5^ peritoneal cavity CD19^+^ B cells isolated by FACs cell sorting from adult CD45.1^+^ mice (12-30 weeks old). *Elavl1^fl/fl^ Mb1^Cre^* and littermate control mice were 8-12 weeks old at the time of B cell transfer. CD45.1^+^ B-1 cell population expansion, relative to host B cell populations expressing CD45.2, was assessed in the peritoneal cavity and in spleen 6 weeks after initial cell transfer.

To assess expansion of self-reactive B-1 cells, 25×10^6^ apoptotic cells were weekly injected (i.v.) into *Elavl1^fl/fl^ Mb1^Cre^* mice and littermate controls. Briefly, thymocytes from C57BL/6 mice were smashed through a 40 µm cell strainer. Red blood cell depleted single cell suspension was the cultured at 37°C in RPMI 1640 Medium (Dutch Modification) supplemented with 10% FCS, antibiotics, 2 mM L-glutamine and 10 µM β-mercaptoethanol. Cells were treated O/N with 10 µM etoposide to induce cell death. Apoptotic cell bodies were then recovered, washed with PBS and resuspended at a concentration of 1.25 x10^8^ cells/ml. 200 ul of the apoptotic cell suspension was i.v. injected at day 0, 7 and 14 prior endpoint analysis of PtC-and PC-reactive B-1 cells and autoantibodies at day 15.

### Flow cytometry

B cell phenotypic characterization was performed mainly by flow cytometry using antibodies listed in ***Suppl. Table 4***. Briefly, cells were first incubated with an Fc Receptor Blocking antibody (clone 2.4G2, BD biosciences) and with Zombie NIR Fixable Viability (BioLegend) for 15 min. at 4°C in PBS containing 2% FCS (FACS buffer). After washing, cell surface receptor labelling was performed by incubating the cells at 4°C for 30 min. with an antibody mix diluted in FACs buffer. Then, cells were extensively washed with FACs buffer prior cell fixation and permeabilization using the BD Cytofix/Cytoperm™ Fixation and Permeabilization Solution for 15 min at 4°C. After cell washing with BD Cytoperm™ washing solution, intracellular proteins were stained O/N at 4°C with specific antibodies diluted in BD Cytoperm™ washing solution containing 1% FCS. After extensive wash, data was collected in a BD Fortessa cytometer and analysed with FlowJo vX. BrdU staining was performed using the FITC BrdU Flow Kit from BD Biosciences as recommended. Alternatively, fixed and permeabilized cells were treated with TurboDNAse (10 U per sample, Thermo Fisher Scientific) for 2 h. at 37°C prior staining of BrdU with a specific antibody (MoBU-1 clone, Thermo Fisher Scientific), O/N at 4°C. Cell viability was assessed by incubating at 37°C up to 10^7^ cells resuspended in complete RPMI medium containing 10 µM of the CaspACE™ FITC-VAD-FMK in Situ Marker (Promega). After 30 min. cells were stained with the Zombie NIR Fixable Viability Dye (BioLegend) prior surface and intracellular protein staining as described above.

### *In-vitro* cell culture

HEK293T cells were cultured in DMEM supplemented with 5% FCS, antibiotics and 2 mM L-glutamine. Peritoneal cavity and splenic B cells were isolated either using MojoSort Mouse CD19 Nanobeads (BioLegend) or by FACs sorting in a FACSAria Fusion cell sorter as indicated below. B cells were then cultured in RPMI 1640 Medium (Dutch Modification) supplemented with 10% FCS, antibiotics, NEAA, 2 mM L-glutamine, 1 mM sodium pryruvate and 10 µM β-mercaptoethanol. In some cases, recombinant human rhBAFF (PeproTech) was added at a concentration of 200 ng/ml. After 24 or 48h, cells were recovered and stained with Dapi for cell survival analysis and/or specific antibodies for quantitation of B220^+^ CD5^-^ CD43^-^ B-2 and B220^-^ CD5^+^ CD43^+^ B-1 cells by flow cytometry.

### BCR-dependent signalling

10^6^ peritoneal cavity cells and 5×10^6^ splenocytes were cultured in 200 µl of serum-free Dutch-modified RPMI-1640 medium for 1 h. at 37°C prior stimulation with an AffiniPure F(ab’)₂ Fragment Goat Anti-Mouse IgM (15 µg/ml, Jackson ImmunoResearch) or with pervanadate (25 µM). After 5 to 15 min. stimulation, cells were rapidly fixed by adding 50 µl of BD PhosFlow Fix Buffer I solution and incubated at room temperature for 15 min. Samples were then washed and permeabilised using BD Phosflow Perm Buffer III for at least 30 min. Samples were pre-treated with Fc Receptor Blocking antibody prior protein staining at 4°C, O/N using specific antibodies to detect B-2 and B-1 cells (***Suppl. Table 4***). Antibodies were diluted in BD Phosflow Perm Buffer III containing 1% FCS to block unspecific binding. After cell wash, data were collected in a BD Fortessa cytometer and analysed with FlowJo vX. Relative quantification to non-stimulated cells incubated at 37°C for the same length of time was performed to correct for possible changes in phosphorylation and protein expression associated to B-cell culture.

Pervanadate was freshly prepared as previously described ^71^. Briefly, 30% H_2_O_2_ was first diluted 100 times with a 20mM HEPES buffer at pH 7.3. 50 µl of this solution was mixed gently with 940 µl of H_2_O and 10 µl of a 100 mM sodium orthovanadate (Na_3_VO_4_) solution. After 5 minutes incubation at room temperature, a small amount of catalase (Sigma Aldrich) was added in with the help of tip. Mix gently by flicking the tube and check for a burst of bubbles indicating enzymatic reaction. Pervanadate is kept at 4°C and discarded after 1 hour. Sodium orthovanadate stock should be colourless. Discard if it shows a yellowish colour and prepare a new stock that can be kept at -20°C.

### Enzyme-linked immunosorbent assay (ELISA)

Total, PC- and PtC-specific antibodies were measured in mouse serum as previously described^72^. A standard ladder was used for quantitation of IgM and IgA levels. On the contrary, relative concentrations of IgM antibodies specific against PC and PtC antigens was calculated by serial serum dilution and endpoint titre calculation. Reagents and antibodies used for capture and detection are listed in ***Suppl. Table 4***.

### Generation and analysis of 3’UTR luciferase reporters

Gene blocks of wild-type and mutant 3’UTR sequences of the *Ighm* mRNA transcripts ENSMUST00000177715 and ENSMUST00000103426, encoding the membrane and soluble protein isoforms respectively, were synthesized by IDT technologies and cloned into the psiCheck2 dual luciferase reporter plasmid (Promega). These plasmid constructs were transfected into HEK293T cells using Lipofectamine 2000 (Thermo Fisher Scientific) used as recommended. Luciferase signal was measured 48h after cell transfection using a Dual-Luciferase® Reporter Assay System from Promega. Relative luciferase units were calculated by dividing renilla luciferase counts by firefly luciferase counts to correct for any differences in transfection efficiency. Later, data was relatively quantified to the control group. At least three independent experiments were performed by triplicate. Pooled data from all experiments is shown in each figure panel. Mann-Whitney tests were performed for statistical analyses.

### RNA extraction and RT-qPCR

Total RNA from HEK293T cells was isolated using Trizol and ethanol precipitation. Abundance of renilla luciferase and firefly luciferase mRNAs was measured by RT-qPCR using 5 ng. of mRNA converted into cDNA with a LunaScript® RT SuperMix Kit (New England Biolabs) and a LightCycler® FastStart DNA Master SYBR Green I kit (Roche) using specific primers previously described ^73^. qPCR in a LightCycler® (Roche) was performed using the following program: incubation at 95°C for 5 min; amplification in 40 cycles of 95°C for 15 sec, 60-62°C for 15 sec and 72°C for 30 sec; melting curve and cooling at 10°C. *Hprt* and/or 18S rRNA were also tested to normalize mRNA abundance. Renilla luciferase mRNA abundance was relative quantified to the levels of firefly luciferase mRNA to account for any differences in cell transfection efficiencies.

### Cell sorting and RNA sequencing

Peritoneal cavity B-1a cells and B-2 cells from *Elavl1^fl/fl^* and *Elavl1^fl/fl^ Mb1^Cre^* mice were FACS-sorted using a FACSAria Fusion cell sorter into complete RPMI medium. Briefly, after cell recovery for the peritoneal cavity, cells were incubated at 4°C in PBS containing 2% FCS, 1 mM EDTA and Fc Receptor blocking antibody. Then, cells were stained as followed: B-2 cells - CD19^+^ B220^hi^ CD43^-^ CD5^-^ CD11b^low^ and B-1a cells - CD19^+^ B220^low^ CD43^+^ CD5^+^ CD11b^hi^.

Total RNA was isolated using a RNeasy Micro Kit from Qiagen and libraries generated using the NEBNext® Single Cell/Low Input RNA Library Prep Kit for Illumina® (New England BioLabs). NEBNext® Multiplex Oligos for Illumina® (Dual Index Primers Set 1 or 2) were used for sample dual indexing and library production. Multiplexed samples were sequenced at BGI Genomics (China) using a DNBSeq platform (100bp paired-end mode).

### Crosslinking Immunoprecipitation and sequencing

Individual nucleotide cross-linking immunoprecipitation (iCLIP) and sequencing was performed to annotate the HuR:RNA interactome as previously described ^32, 74, 75^. Briefly, peritoneal cavity cells were isolated from a minimum of 8 C57BL/6 mice (8-12 weeks old) using ice-cold PBS supplemented with 2% FCS and right after irradiated with UV light (600 mJ/cm2 using a Stratalinker 2400) to preserve *in-vivo* protein:RNA interactions. Then, CD19^+^ B cells were positively isolated using MojoSort Mouse CD19 Nanobeads (BioLegend). 1-2×10^7^ B cells were lysed for 15 min. using 1 ml of ice-cold RIPA buffer (50 mM Tris-HCl - pH 7.4, 100 mM NaCl, 1% IGEPAL CA-630, 0.1% SDS, 0.5% Sodium deoxycholate, 10 units/ml of RNase Out and protease inhibitor cocktail at a 1 in 100 dilution (Sigma Aldrich)). Then, samples were sonicated in 3 cycles of 10 s. each, and centrifuged at 15000 rpm at 4°C. Supernatant was transferred to a new ice-cold eppendorf tube prior adding TurboDNase (10 U/ml of lysate, ThermoFisher Scientific) and RNase I (0.167 U/ml, ThermoFisher Scientific). Samples were incubated exactly 3 min. at 37°C shaking at 1100 rpm for genomic DNA and RNA digestion. Immunoprecipitation of HuR-RNA complexes was performed using 3 µg. of HuR antibody (clone 3A2, Santa Cruz Bio.) as previously described ^40^. Alternatively, as negative control, we used a mouse IgG1 isotype control antibody which failed to precipitate any RNA and to produce an iCLIP library. After O/N incubation at 4°C, beads were washed 3 times with a high-stringent washing buffer (50 mM Tris-HCl pH 7.4, 1 M NaCl, 1 mM EDTA, 1% NP-40, 0.1% SDS and 0.5% sodium deoxycholate) and once with PNK washing buffer (20 mM Tris-HCl pH 7.4, 10 mM MgCl2, 0.2% Tween-20) prior linking a L3-ATT-App adaptor sequence at the 3’ end of the RNA (L3-ATT-App DNA Linker (/5rApp/WN ATT AGA TCG GAA GAG CGG TTC AG/3Bio/)). First, 3’end RNA dephosphorylation was carried out by combining 0.5 U of FastAP alkaline phosphatase (Thermo Fisher Scientific) and 10 U of PNK (New England Biolabs). Then, after stringent washes, beads were resuspended with 25 µl of the following ligation mix: 3 μl 10x ligation buffer (500 mM Tris-HCL pH 7.5 and 100 mM MgCl2), 0.8 μl 100% DMSO, 75 U of T4 RNA ligase I (New England BioLabs), 15 U of RNase Out (Thermo Fisher Scientific), 5 U of PNK (New England BioLabs), 11.25 μl of 40% PEG8000 and 2.5 ul of 10 μM L3-ATT-App adaptor sequence. Extensive washes were performed after IP using high-stringent washing buffer and PNK washing buffer. One tenth of the sample was alternatively ligated to a pre-adenylated infrared L3-IR-App adaptor (/5rApp/AG ATC GGA AGA GCG GTT CAG AAA AAA AAA AAA /iAzideN/AA AAA AAA AAA A/3Bio/ coupled with IRdye-800CW-DBCO (LI-COR)) for HuR:RNA complex visualization by SDS-Page electrophoresis using 12-15% Novex™ TBE-Urea pre-cast gels and protein:RNA complex transfer to nitrocellulose membranes. Protein:RNA complexes above 20-KDa the molecular weight of HuR protein, were isolated by incubating specific nitrocellulose membrane fragments with PK buffer (100 mM This-Cl pH 7.5, 100 mM NaCl, 1 mM EDTA, 0.2 % SDS and 20 U of proteinase K [Roche]). After 60 min. at 50°C, RNA was isolated using phenol/chloroform extraction and ethanol precipitation. irCLIP_ddRT_primer 43 (/5Phos/WWW AGTTA NNNN AGG CAG GAA GAG CGT CGT GAT/iSp18/GGA TCC/iSp18/TAC TGA ACC GC) was for RNA to cDNA conversion using SuperScript IV reverse transcriptase (Thermo Fisher Scientific). cDNA was purified with Agencourt AMPure XP beads (Beckman), circularised with CircLigase II (Epicentre) and PCR amplified using Solexa P5/P7 primers. iCLIP library sequencing service (100bp, single-end sequencing) was provided by BGI Genomics, China.

### Bioinformatics

Raw reads from different sequencing lanes were concatenated using bash cat utility and trimmed using flexbar (v.3.5.0) to remove NEBNext® adapters as recommended by the manufacturer (with parameters --adapter-trim-end LEFT, --adapter-revcomp ON, --adapter- revcomp-end RIGHT, --htrim-left GT, --htrim-right CA, --htrim-min-length 3, --htrim-max- length 5, --htrim-max-first, --htrim-adapter, --min-read-length 2 for the first flexbar command and –interleaved, --adapter-trim-end RIGHT, --min-read-length 2 for the second flexbar command). mRNAseq data was further processed using the nextflow nf-core/RNAseq pipeline (v3.8.1) ^76^. Briefly, reads were trimmed with Cutadapt (v3.4) and Trimgalore (v.0.6.7). FASTQC-0.11.9 was used to assess the quality of resulting reads. Paired-end reads were aligned to the mouse genome (GRCm39-v106) and quantified with STAR-2.7.10a (with options --runMode alignReads --outSAMtype BAM SortedByCoordinate --quantMode TranscriptomeSAM GeneCounts --twopassMode Basic, with option --sjdbOverhang 100). Alignment files were indexed using SamTools (v. 1.15.1). mRNAseq data analysis was performed using R / R Studio by importing salmon generated counts for transcripts collapsed to gene level using *tximport* R package ^77^. DESeq2 (v1.28.0) ^78^ was used for differential expression analysis using default parameters. P values were adjusted using the Benjamini and Hochberg correction. Significant changes in gene expression were defined as those genes with a p adjusted (padj) value below 0.05. Analysis of alternative splicing was performed with rMATS-4.1.2 and Miniconda3 (using options--readLength 150 --libType fr-unstranded and Ensembl GRCm39-v103 mouse genome annotation). Mouse annotation was the same as the one used for STAR indexing and alignment. Differentially splicing events were considered significant if the false discovery rate (FDR) was below 0.05 and the absolute inclusion level was higher than 0.1 in a scale from 0 to 1. Gene ontology enrichment analyses were performed using Enrichr (10.1186/1471-2105-14-128 and 10.1002/cpz1.90) using default settings. iCLIP analyses were performed using Flow (https://app.flow.bio/) which integrates nf-core/clipseq v1.0 for iCLIP sequencing data analyses (https://github.com/nf-core/clipseq). Default parameters were applied to retrieve single-nucleotide resolution binding sites, binding scores, genomic location and genomic features.

### Statistics and reproducibility

Statistics were performed in R or using Prism 7 or later. The statistical tests used in all cases are indicated at the bottom of each figure legend. Briefly, we used Mann-Whitney or t-tests for pairwise comparisons. ANOVA followed by Tukey’s multiple comparison tests were used when having more than 2 groups. Two-sides analyses were performed in all cases. Kolmogorov–Smirnov test allowed assessing equality of one-dimensional probability cumulative distributions. Benjamini and Hochberg test was used with transcriptomics data for multiple testing and p adjusted value calculation. All experiments were performed at least twice. In most cases data was reproduced in more than three independent experiments.

### Data availability

RNAseq and iCLIP raw data can be accessed at Gene Expression Omnibus (GEO) using the accession numbers GSE276801 and GSE276641. The rest of raw and processed data is readily available under request.

## SUPPLEMENTAL INFORMATION

**Supplementary figure 1.**
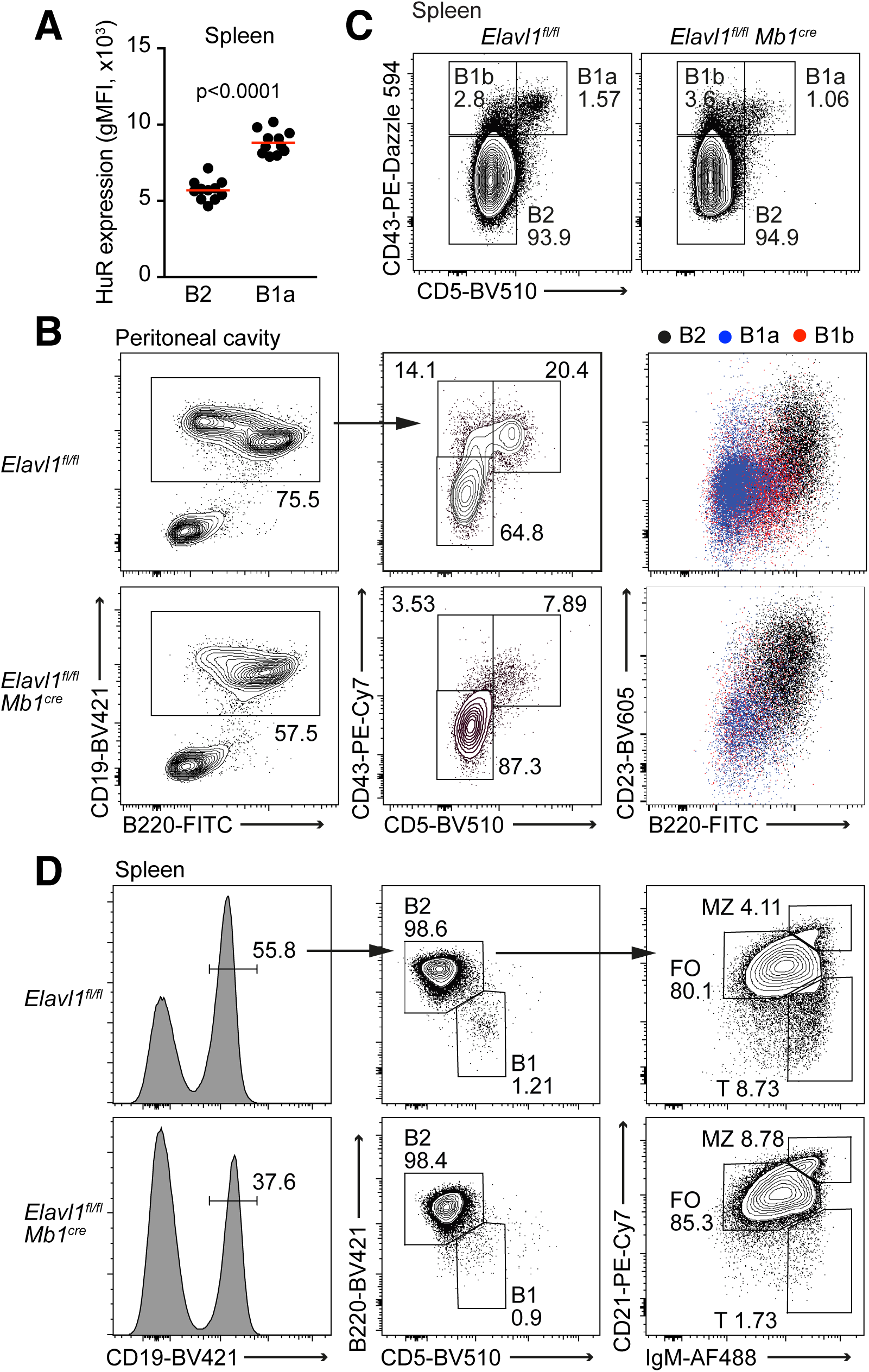
Characterization of B-1 and B-2 cells in peritoneal cavity and spleen. **A,** HuR expression in splenic B-1 and B-2 cells measured by flow cytometry. Data pooled from two out of more than three independent experiments with a minimum of 3 mice. **B,** Counter plots showing the flow cytometry gating strategy to identify B-1a, B-1b and B-2 cells in the peritoneal cavity of control and *Elavl1^fl/fl^ Mb1^Cre^*mice. Right panels, superposed dot plots showing the expression of CD23 and B220 in the different B cell subsets. **C,** Counter plots showing the percentage of splenic B-1a, B-1b and B-2 cells in control and *Elavl1^fl/fl^ Mb1^Cre^* mice. Previously gated on CD19+ cells. **D,** Flow cytometry gating strategy to analyse all B cell subsets in the spleen. All data is representative of more than three independent experiments performed with at least 3 mice per genotype.

**Supplementary figure 2.**
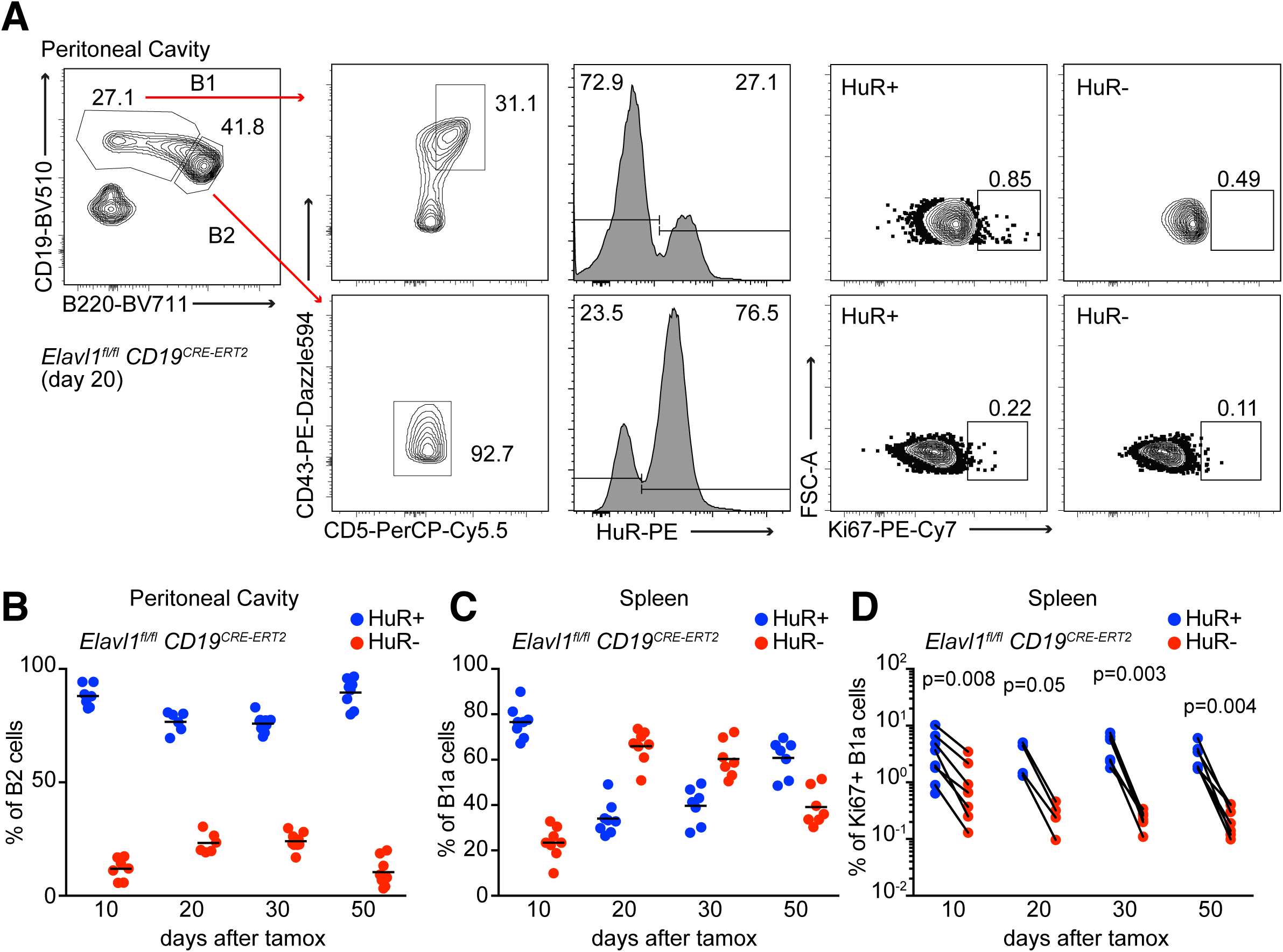
Analysis of tamoxifen-induced deletion of HuR in B cells from *Elavl1^fl/fl^ CD19^CRE-ERT^*^2^ mice. **A**, Flow cytometry gating strategy for the analysis of B cell population in control and *Elavl1^fl/fl^ CD19^CRE-ERT^*^2^ mice after tamoxifen-induced deletion of HuR. **B,** Proportion of HuR^+^ and HuR^-^ B-2 cells in the peritoneal cavity of the *Elavl1^fl/fl^ CD19^CRE-ERT^*^2^ mice shown in Fig. 3C. **C,** Analysis of HuR deletion in splenic B-1a cells from *Elavl1^fl/fl^ CD19^CRE-ERT^*^2^ mice at the indicated endpoints. **D,** Comparative analysis of the proportion of HuR^+^ Ki67^+^ and HuR^-^ Ki67^+^ splenic B-1a cells in *Elavl1^fl/fl^ CD19^CRE-ERT^*^2^ mice.

**Supplementary figure 3.**
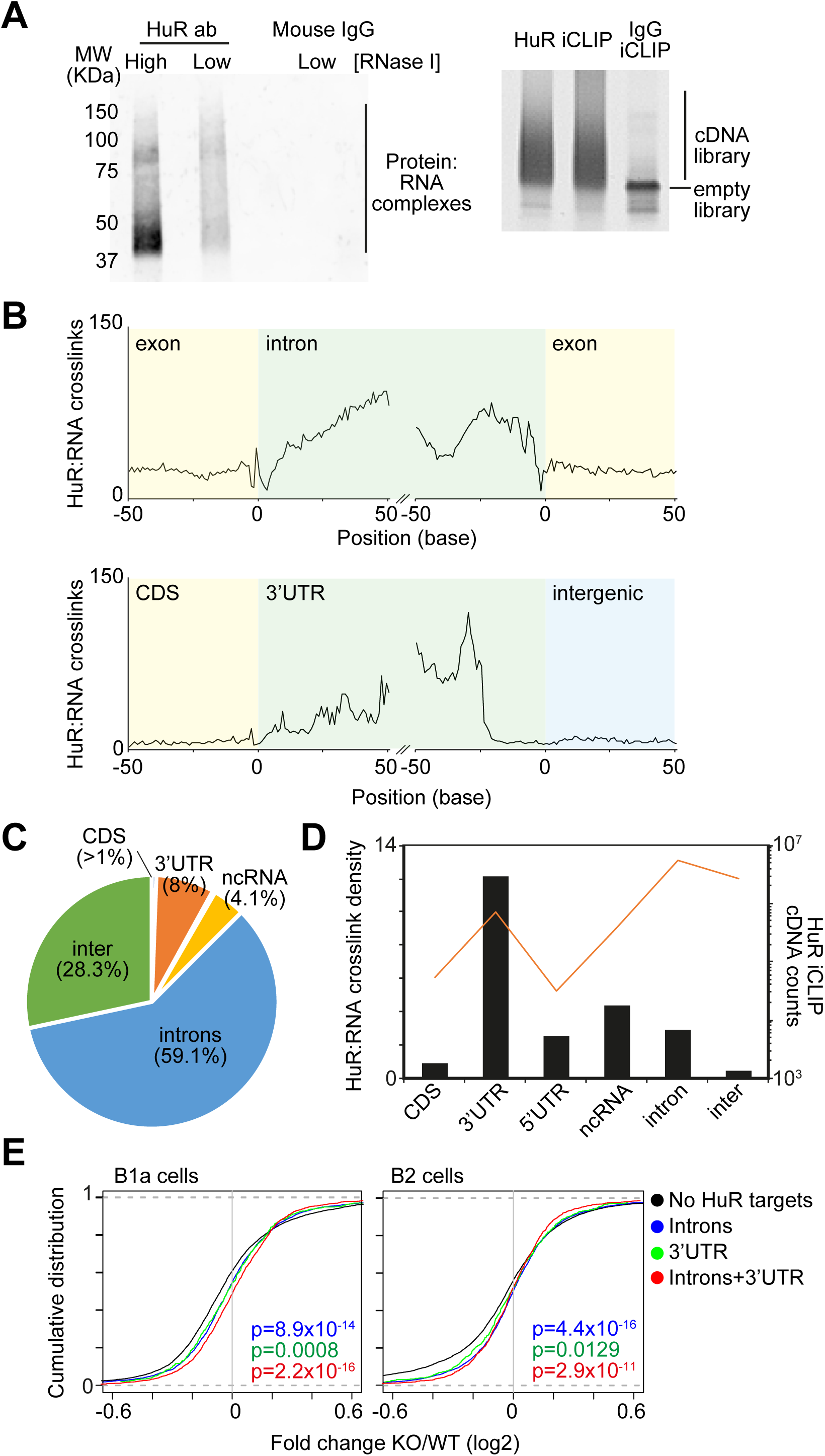
Characterization of the HuR:RNA interactome in peritoneal B cells. **A,** Detection of the HuR:RNA complexes immunoprecipitated using an anti-HuR or an isotype antibody and separated by SDS-Page and transfer into a nitrocellulose membrane. Right, visualization of HuR iCLIP libraries generated with the immunoprecipitated HuR:RNA complexes shown in the left. **B,** RNA maps showing the density of HuR:RNA crosslinks annotated in the exon-intron, intron-exon, CDS-3’UTR and 3’UTR-intergenic boundaries. **C,** Pie chart indicated the percentage of HuR:RNA crosslinks annotated to each of the genomic features indicated. **D,** Quantitation of the HuR binding density and the number of unique cDNA counts mapped to each genomic feature. **E,** Cumulative distribution plots showing gene expression changes in HuR KO B-2 and B-1a cells for those genes targeted or not by HuR at introns, 3’UTRs or both.

**Supplementary figure 4.**
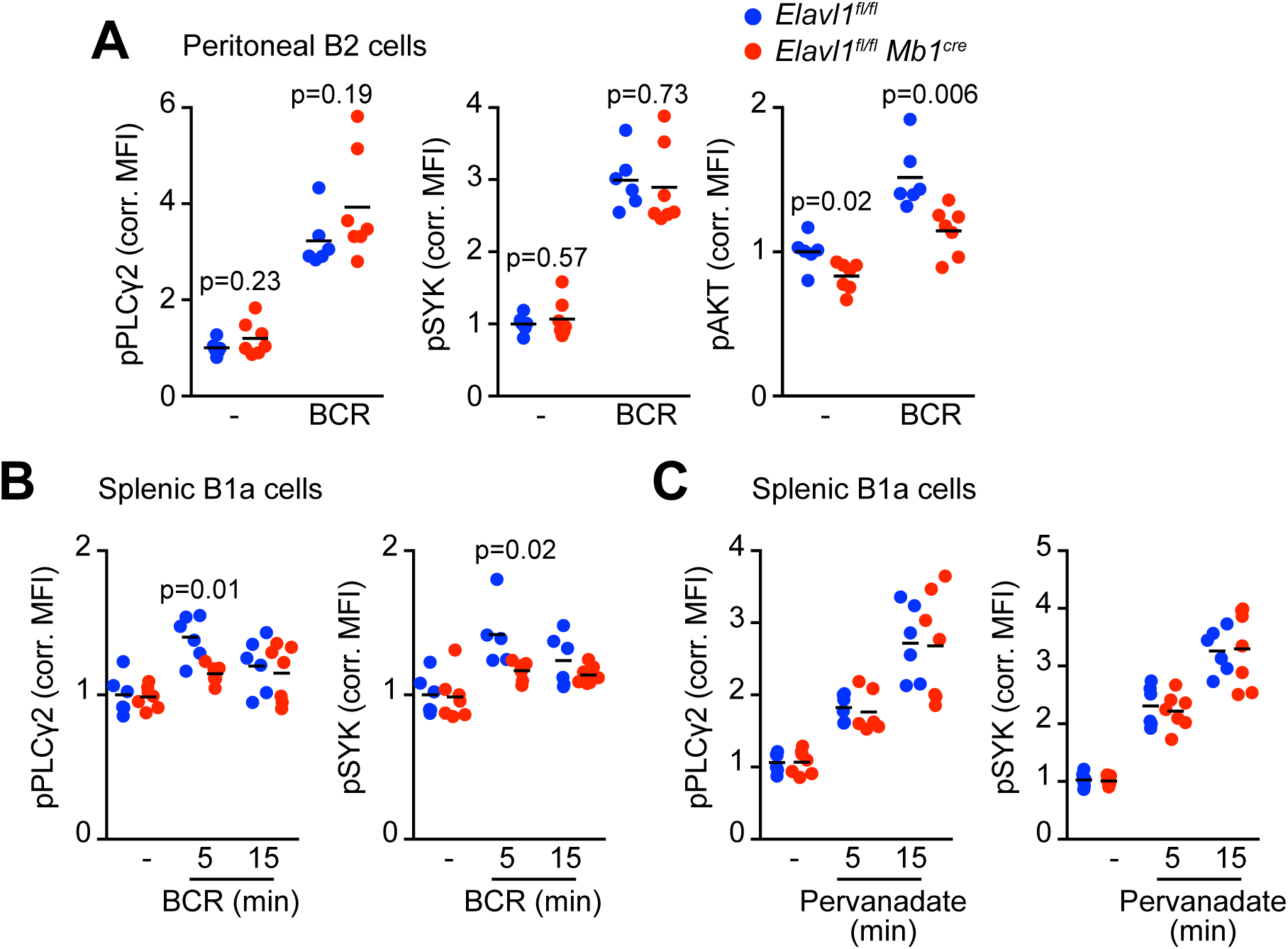
Analysis of BCR-dependent signalling activation in HuR KO B-1 and B-2 cells. **A,** Quantitation of pPLCγ2, pSYK and pAKT in peritoneal B-2 cells after BCR-mediated activation for 5 min. **B, C,** Analysis of PLCγ2 and SYK phosphorylation in splenic B-1a cells stimulated *in-vitro* with an anti-IgM (**B**) or with pervanadate (**C**) for 5-or 15-min. Data shown in this figure were pooled from the same mice shown in Fig. 5B, 5C and 5D. Data shown in all panels was normalised to the non-treated B cells from control mice. Two-tailed Mann-Whitney tests were performed for statistical analysis.

**Supplementary figure 5.**
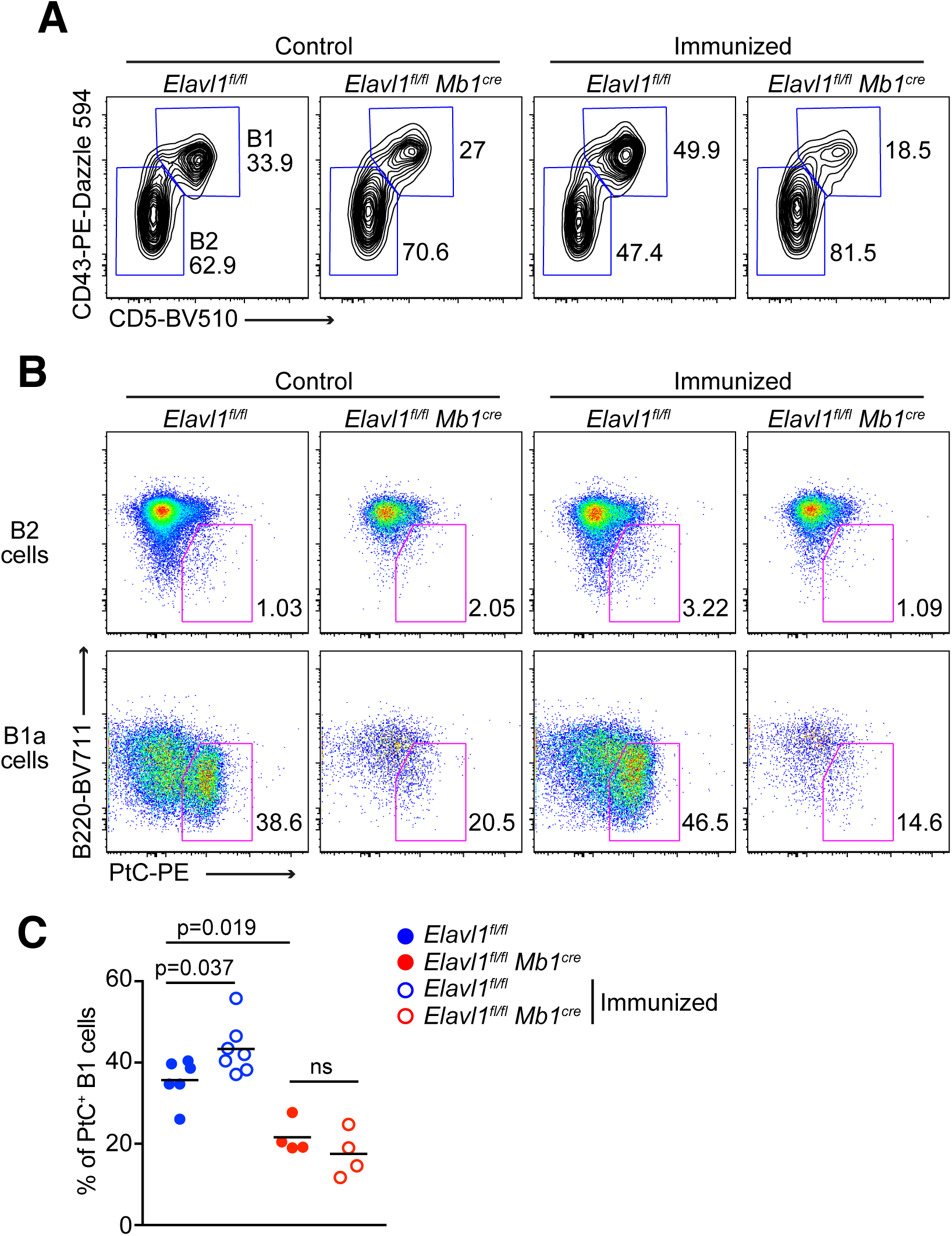
Analysis of self-antigen B-1 cell expansion upon T-independent type 2 immunization. **A,** Representative contour plots for the detection of peritoneal B-1 and B-2 cells in control and *Elavl1^fl/fl^ Mb1^Cre^*mice upon immunization with apoptotic bodies. **B,** Pseudo-colour dot plots showing the proportion of PtC-specific B cells within the B-2 and B-1 compartments as defined in A. **C,** Percentage of peritoneal B-1 cells that recognise the self-antigen PtC in control and *Elavl1^fl/fl^ Mb1^Cre^* mice upon immunization with apoptotic bodies shown in Fig 7C. Two-tailed Mann-Whitney tests were performed for statistical analysis.

**Supplementary table 1. Differential gene expression analyses comparing control and HuR KO B-1a and B-2 cells.**

**Supplementary table 2. HuR:RNA interactome analysis from peritoneal B cells.**

**Supplementary table 3. Pathway enrichment analyses.**

**Supplementary table 4. List of antibodies.**

## Notes

### Competing Interest Statement

The authors have declared no competing interest.

